# Lipid-mediated insertion of Toll-like receptor (TLR) ligands for facile immune cell engineering

**DOI:** 10.1101/840876

**Authors:** Michael H. Zhang, Emily M. Slaby, Georgina Stephanie, Chunsong Yu, Darcy M. Watts, Haipeng Liu, Gregory L. Szeto

## Abstract

Cell-based immunotherapies have tremendous potential to treat many diseases, such as activating immunity in cancer or suppressing it in autoimmune diseases. Most cell-based cancer immunotherapies in the clinic provide adjuvant signals through genetic engineering to enhance T cell functions. However, genetically encoded signals have minimal control over dosing and persist for the life of a cell lineage. These properties make it difficult to balance increasing therapeutic efficacy with reducing toxicities. Here, we demonstrated the potential of phospholipid-coupled ligands as a non-genetic system for immune cell engineering. This system provides simple, controlled, non-genetic adjuvant delivery to immune cells via lipid-mediated insertion into plasma membranes. Lipid-mediated insertion (depoting) successfully delivered Toll-like receptor (TLR) ligands intracellularly and onto cell surfaces of diverse immune cells. These ligands depoted into immune cells in a dose-controlled fashion and did not compete during multiplex pairwise loading. Immune cell activation could be enhanced by autocrine and paracrine mechanisms depending on the biology of the TLR ligand tested. We determined that depoted ligands can functionally persist on plasma membranes for up to four days in naïve and activated T cells, enhancing their activation, proliferation, and skewing cytokine secretion. Depoted ligands provide a persistent yet non-permanent adjuvant signal to immune cells that may minimize the intensity and duration of toxicities compared to permanent genetic delivery. Altogether, these findings demonstrate potential for lipid-mediated insertion (depoting) as a universal cell engineering approach with unique, complementary advantages to other cell engineering methods.

## Introduction

Advances in drug delivery have enhanced our understanding of basic biology and generated novel therapies. In cell-based immunotherapy, drugs are administered systemically to target immune cells *in vivo*, or carried by *ex* vivo-primed autologous immune cells that are reinfused into patients. Strategies to engineer cells as drug-carriers *ex vivo* include genome editing, conjugating biomaterial-based carriers to cell surfaces, and binding of the plasma membrane for passive diffusion of immunostimulatory ligands (1, 2). Despite successful implementation in the clinic, cell-based immunotherapies still face challenges of implementing simple and efficient delivery that improve therapeutic responses. For example, genome editing can decrease cell viability and has low efficiency in some cell types of interest, particularly T cells (3–6). Some immunostimulatory ligands, such as small-molecule drugs, cannot be genetically encoded. Biomaterials can be assembled into carriers with defined size, shape, cargo loading, composition, and physiochemical parameters that enable controlled delivery to intracellular cell compartments for autocrine signaling or to surrounding cell surface receptors for paracrine signaling (2). However, optimization of nanoparticle design parameters is complex, and must be either universally compatible or specifically designed for *ex vivo* anchoring to and uptake by different immune cells (7). Here, we propose a simplified delivery platform that couples diverse biomolecular cargo to phospholipids that directly insert into plasma membranes for universal loading into plasma membranes of cells without the need for gene editing or complex biomaterial design.

Biomolecules have been previously conjugated to phospholipids for cell loading by plasma membrane interactions. Our previous study generated potent vaccines by exploiting the affinity of lipids to bind albumin and deliver antigen and adjuvant conjugates to lymphoid organs (8). *Ex vivo* modification with lipid conjugated immunostimulatory ligands has also been investigated for some adjuvants. In cancer, lipid-mediated delivery of T-cell adjuvants elicit tumor regression in multiple preclinical models of cancer (9, 10). Lipid conjugated adjuvants also strongly associate with plasma membranes of dendritic cells and tumor cells (11, 12). However, previous studies have not fully characterized the efficiency of lipid-mediated insertion into plasma membranes, or expanded this delivery approach to diverse immunostimulatory cargoes.

Toll-like receptors (TLRs) are one family of molecules that are heavily used to enhance immune responses. TLRs are a major contributor to innate immune sensing (4), and also directly implicated in adaptive immunity during many diseases (13–16). For example, TLR2 is a cell surface receptor that senses molecules from microbial cell-walls. TLR2 activation results in increased pro-inflammatory cytokine secretion, suppression of regulatory T cells, and enhanced sensitivity of cytotoxic T cells (3, 14, 17). TLR9 is an intracellular receptor that senses unmethylated CpG DNA from viruses and bacteria. TLR9 ligands are frequently used as vaccine adjuvants (18, 19), and can directly enhance proliferation and survival of T cells (20, 21). Successful delivery of both TLR2 and TLR9 ligands has demonstrated promising therapeutic responses, particularly in cancer (14, 22–24). However, improved delivery approaches are needed to more easily deliver cell surface and intracellular ligands to diverse immune cells.

We report that lipid-conjugated TLR2 and TLR9 ligands can rapidly and simply insert into immune cell plasma membranes (hereafter termed depoting), delivering ligands to both cell surface and intracellular receptors. We analyzed ligand loading, dynamics of ligand persistence, and activation of mouse immune cells to demonstrate the feasibility of depoting to provide paracrine and autocrine signals that can enhance immune cell function. This study provides proof-of-concept that depoting immunostimulatory ligands into plasma membranes can enhance cell function, detail key features of this platform including dynamics of single and multiplex ligand loading and turnover, and provide a new method for engineering cell-based therapies that can complement any existing methods.

## Methods

### TLR2 and TLR9 Ligands

Synthetic ligands for TLR2, Pam_2_CSK_4_ and Pam_3_CSK_4_, and their biotinylated variants were purchased from Tocris Bioscience and InvivoGen. TLR9 ligand, CpG oligonucleotide 1826 (5’-tccatgacgttcctgacgtt-3’ with a phosphorothioated backbone) (CpG) and fluorescein (FAM)-labeled CpG, were commercially synthesized (Integrated DNA Technologies). Diacyl stearoyl (C18) lipid conjugated CpG (lipid-CpG) and FAM-labeled lipid-CpG were made as previously described by synthesizing diacyl C18 lipid phosphoramidite and conjugating to either CpG or CpG-FAM on a ABI 394 synthesizer on 1.0 micromole scale (12). Lipid-CpG was purified by reverse phase HPLC with a C4 column (BioBasic4, 200 mm × 4.6 mm, Thermo Scientific). A gradient eluent (Sigma-Aldrich) was implemented with 100 mM triethylamine-acetic acid buffer (pH 7.5) and acetonitrile (0–30 min, 10–100%).

### Isolation of Naïve and Primed Mouse Immune Cells

All procedures with animals and animal-derived materials were approved by the UMBC Institutional Animal Care and Use Committee (OLAW Animal Welfare Assurance D16-00462). C57BL/6 mice and CD45.1+ (B6.SJL-Ptprca Pepcb/Boy) congenic mice from Jackson Laboratory were bred in the UMBC animal facility and used for all experiments. Spleens from 12-52 week old C57BL/6 mice were mashed through a 40-μm cell strainer treated with ACK lysis buffer (1 mL per spleen, ThermoFisher) for 5 min at 25°C to lyse red blood cells. Naive CD4+ and CD8+ T cells were isolated using a negative selection cocktail containing the following biotinylated mouse antibodies (Biolegend): TCR γ/δ (clone GL3), CD24 (clone M1/69), TER-119 (clone TER-119), CD49b (clone HMα2), CD45R/B220 (clone RA3-6B2), CD19 (clone 6D5), CD11c (clone N418), and CD11b (clone M1/70). B cells were isolated from splenocytes using a negative selection cocktail containing the following biotinylated mouse antibodies: CD43 (clone 1B11), CD90.2 (clone 30-H12), Gr-1 (clone RB6-8C5), TER-119 (clone TER-119), CD49b (clone HMα2), CD11b (clone M1/70), CD8 (clone 53-6.7), and CD4 (clone H129.19). Antibody-bound cells were depleted with Rapidsphere streptavidin magnetic beads according to the manufacturer’s instructions (STEMCELL Technologies).

Primed T cells were obtained by culturing splenocytes in complete RPMI 1640 media supplemented with 10% fetal bovine serum (FBS; ThermoFisher), concanavalin A (2 μg/mL; Sigma-Aldrich), and IL-7 (2 ng/mL, Biolegend) at 37°C for 2 days. Ficoll-Pague Plus (GE Healthcare Life Sciences) gradient separation was used for dead cell removal by centrifugation at 500 x g for 20 min with no break. Primed T cells were isolated with the negative selection cocktail described above.

### Lipid-Mediated Insertion (Depoting) of Lipid-Conjugated Ligands

Unless specified otherwise (in Fig. 2), splenocytes, isolated B cells, or isolated T cells were incubated with CpG (5 μM), lipid-CpG (5 μM), Pam2CSK4 (10 μg/mL), or Pam3CSK4 (10 μg/mL) for 1 hour in complete RPMI 1640 media supplemented with 10% fetal bovine serum. Splenocytes or T cells were then washed 3 times with PBS plus 1% bovine serum albumin to remove unbound ligand.

Cells were incubated with αCD16/32 antibody (clone 93; Biolegend) to block non-specific antibody binding by Fc receptors for 5 min at 25°C. Cells were stained with the following antibodies: CD4 (clone GK1.5; PerCP/Cyanine5.5), CD8a (clone 53-6.7; APC), and B220 (clone RA3-6B2; PE/Cy7) 15 min at 25°C. PE-conjugated streptavidin was used to bind depoted biotinylated Pam2CSK4. Cell viability was determined by LIVE/DEAD™ exclusion staining per manufacturer’s instructions (ThermoFisher).

### TLR2 Ligand Persistence and Bystander Cell Activation Assays

Naive or primed T cells were rested in complete RPMI 1640 media and supplemented with 10% FBS after depoting with Pam_2_CSK_4_ or Pam_3_CSK_4_ as described above. At selected time-points (0, 1, 2, 5, or 8 days post-depoting), T cells were washed and fixed with 4% paraformaldehyde (Sigma) for 15 min at 25°C. After washing 5 times with PBS + 1% BSA, at 1000 x *g* for 5 min, T cells were cultured in complete RPMI 1640 media and supplemented with 10% FBS. Bystander B cells were added in co-cultures at a 1:1 B cell:T cell ratio. Naive T cells without fixation were immediately cultured with bystander B cells. After 2 days of incubation at 37°C and 5% CO_2_, B-cell activation was measured by fluorescent staining for MHCII (clone M5/114.15.2; FITC) and CD69 (clone H1.2F3; PerCP/Cy5.5).

### Lipid-Conjugated TLR9 Ligand APC Activation Assay

Isolated B cells from C57/BL6 mice were cultured in complete RPMI 1640 media supplemented with 10% FBS after depoting with lipid-CpG and combined with an equal number (50,000 cells) of isolated CD45.1^+^ B cells. After 2 days of co-culture at 37°C and 5% CO_2_, B cells were identified by B220 (clone RA3-6B2; PE-Cy7) CD45.1 (clone A20; PE) antibody, and B-cell activation was measured by MHCII (clone M5/114.15.2; FITC) and CD69 (clone H1.2F3; PerCP/Cy5.5).

### T-cell Activation Assays

Naïve T cells were labeled with 5 μM of carboxyfluorescein succinimidyl ester (CFSE, ThermoFisher). Labeled cells were cultured in RPMI 1640 media and supplemented with 10% FBS as well as αCD3/CD28 coated Dynabeads™ (Thermo) at a 1:5 bead to T cell ratio after depoting of lipid-CpG, Pam2CSK4, and/or Pam3CSK4 ligands. Non-depoted T cells were treated with soluble lipid-CpG (5 μM), Pam2CSK4 (10 μg/mL), and/or Pam3CSK4 (10 μg/mL). Cell proliferation was measured by CFSE dilution after 3 days, and cell activation was measured by fluorescent staining for CD25 (clone PC61; PE). Cell proliferation index and division index were calculated using FlowJo LLC software. Proliferation index is defined as the total number of cell divisions divided by the number of divided cells, whereas division index is the total number of cell divisions divided by the number of total original cells (25). Paracrine-enhanced proliferation of bystander T cells was measured as described above, with αCD3/CD28 beads added at a 2:5 bead to T-cell ratio. T-cell supernatents were collected on day 2 for IL-2, IL-4, and IFNγ detection by ELISA per manufacturer’s instructions (BioLegend).

### Flow Cytometry and Microscopy Analyses

Fluorescently labeled cells were analyzed on a BD LSRII or Beckman Coulter CyAn ADP flow cytometer. The LSRII flow cytometer consisted of 405 nm, 488 nm, 561 nm, and 640 nm excitation laser lines. The CyAn ADP flow cytometer consists of 405 nm, 488 nm, and 635 nm excitation laser lines. Data were analyzed using FlowJo LLC software v10 (Tree Star Inc.). Cells for confocal analysis were blocked with normal goat serum and labeled with early endosome marker 1 (EEA1) (clone C45B10; Cell Signaling Technologies) for 12 hours at 4°C. Cells were fluorescently labeled with AF647 anti-rabbit IgG (Invitrogen) and DAPI for 60 min at 25°C. Confocal anaylsis was performed on a TCS SP5 microscope (Leica Microsystems). The excitation laser lines used were 405 nm, 488 nm, and 633 nm.

### Enzyme-Linked Immunosorbent Assay (ELISA) for Lipid-Conjugated Ligand Detection

One million purified T cells were depoted with FAM-labeled lipid-CpG, biotinylated Pam2CSK4, or biotinylated Pam3CSK4 as described above, and then lysed with Glo lysis buffer (Promega) for 10 min at 25°C. Lysates were collected and plated onto Nunc MaxiSorp ELISA plates (Thermo). After 2 hours of incubation at 25°C, FAM-labeled lipid-CpG was excited at 488 nm, and fluorescent emission at 520 nm was quantified by a microplate fluorescence reader. For Pam2CSK4 and Pam3CSK4 samples, horseradish peroxidase (HRP)-labeled streptavidin was added for 2-hour incubation at 25°C, and then TMB substrate (Thermo) was added to observe changes in absorbance at 450 nm with a microplate spectrophotometer (BioTek).

### Regression and Statistical Analyses

For regression models, a least-squares two-phase decay model was fit to the median fluorescence intensity (MFI) data from the TLR2 ligand persistence assay. Effective half-life, the fast half-life of the two-phase decay, is computed as followed: t1/2 = ln(2/(fast rate constant)).

One-way analyses of variance (ANOVAs) were performed followed by comparisons using Dunnett’s, Tukey’s, Sidak’s, or Holm-Sidak’s correction methods. Comparisons to a hypothetical mean were performed using one-tailed ratio-paired t-tests. All statistical analysis and regression models were performed with Prism software (GraphPad). * p < 0.05; ns, not significant.

## Results

### Lipid-conjugated TLR ligands efficiently inserted into plasma membranes

The ability of lipids to insert into plasma membranes can be exploited for stable anchoring of lipid-conjugated immunostimulatory TLR ligands into cells. This phenomenon is similar to the natural insertion of GPI-anchored proteins. We have previously conjugated the intracellular TLR9 ligand CpG DNA to a diacyl stearoyl (C18) lipid, termed lipid-CpG, and demonstrated strong association with plasma membranes of cancer cells (11). Here, we tested whether lipid-CpG can also depot into plasma membranes of immune cells. Lipid-CpG incubated for 1 hour with mouse splenocytes showed increased cell insertion compared to unmodified CpG (**Fig. 1A**). Interestingly, resting lymphocytes (B cells, CD4^+^ T cells, CD8^+^ T cells) showed increased lipid-CpG insertion. Analysis by confocal microscopy suggested the diacyl lipid domain mediated ligand anchoring with plasma membranes of immune cells, and that ligand uptake was only partially through endosomal pathways as determined by colocalization (**Fig. 1B**). These data demonstrated lipid-mediated cell loading can overcome the challenges of low non-specific uptake by resting lymphocytes (26). We termed this lipid-mediated insertion “depoting,” reflecting the likely rapid partitioning of lipid moieties into the lipid bilayer of plasma membranes (27, 28).

**Fig. 1:**
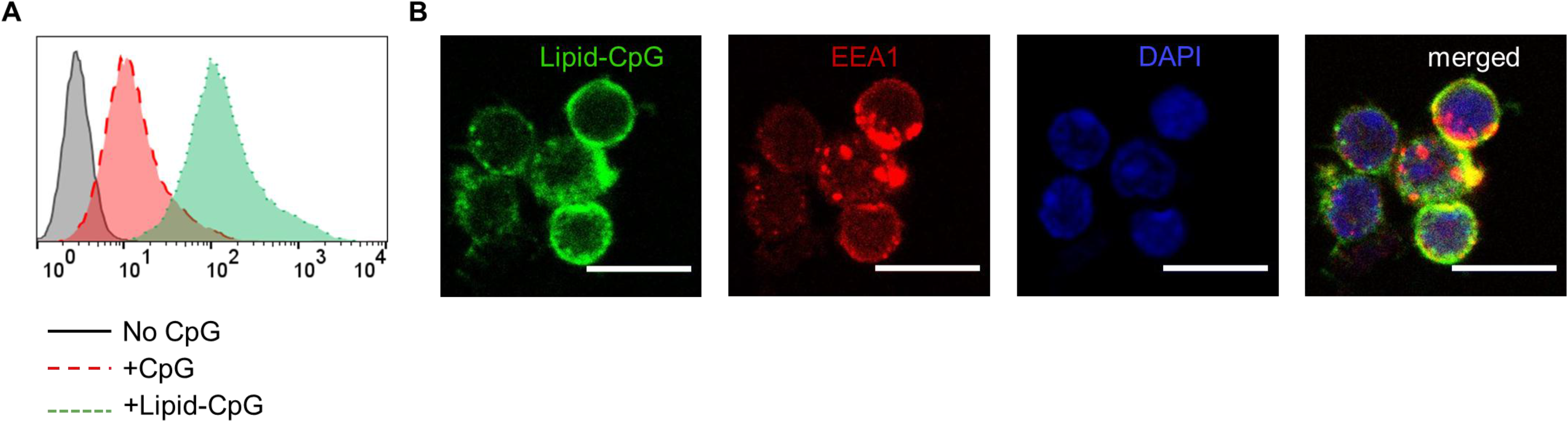
Insertion of TLR9 ligand onto murine immune cells is enhanced by lipid tail. Fluorescein (FAM) labeled CpG DNA or diacyl lipid conjugated CpG (lipid-CpG) were incubated with splenic immune cells at 5 μM for 1 hour at 37°C in supplemented RPMI media. **A)** Fluorescent intensity was measured by flow cytometry. **B)** Confocal micrographs show lipid-CpG coating cell plasma membranes and partial colocalizing with early endosome marker-1 (EEA1). Lipid-CpG (green), EEA1 (red), DAPI (blue). Scale bar = 10 μm.

We next investigated whether a cell surface TLR ligand, Pam_2_CSK_4_, could also be depoted into splenic immune cells given differences in its diacyl palmitoyl (C16) lipid tail. We also tested depoting of Pam_2_CSK_4_ in parallel with lipid-CpG to determine if multiple lipid-tailed ligands can be depoted into cells without competition. Depoting lipid-CpG and Pam_2_CSK_4_ together did not decrease ligand levels compared to single ligand depoting in B220^+^ B cells (**Fig. 2A**), CD4^+^ T cells (**Fig. 2B**), or CD8^+^ T cells (**Fig. 2C**). These data suggested that delivery of TLR2 and TLR9 ligands can be mediated by C16 or C18 lipid tails, respectively, and that depoting 2 ligands was feasible without loading saturation. Further, these data indicated the capacity of plasma membranes for depoted molecules was not saturated under any of our tested conditions. Depoting had no effect on cell viability compared to untreated cells (**Fig. S1**). Overall, our data showed that B and T cells have a high capacity for rapid depoting with multiple lipid-conjugated TLR ligands, and that depoting achieved delivery of ligands into plasma membranes without toxicity to cells.

**Fig. 2:**
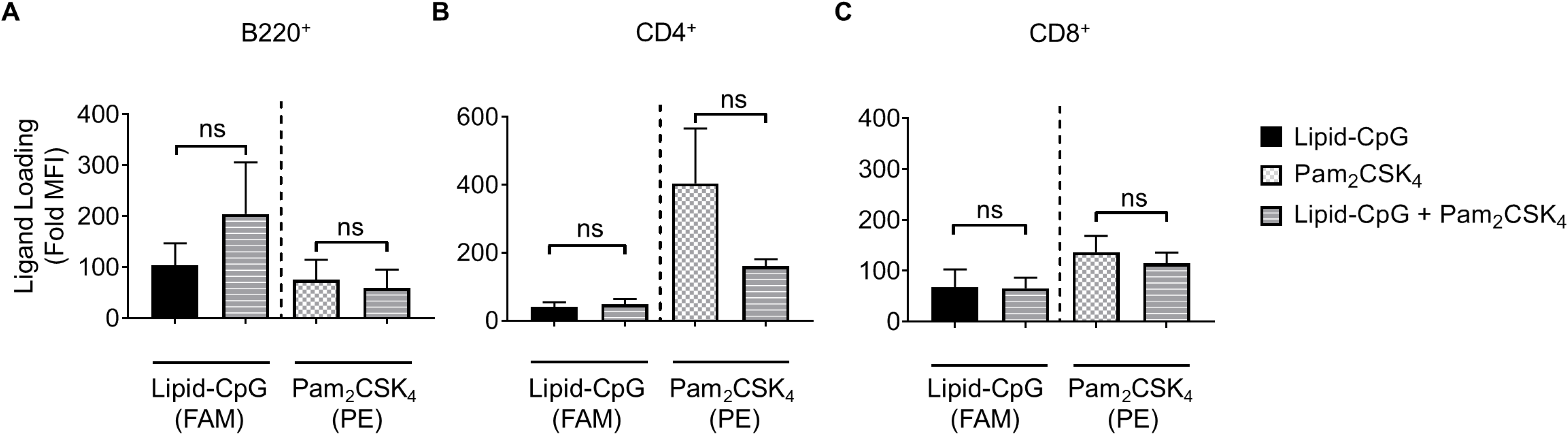
Lipid-tailed ligands efficiently depoted alone and together into resting B and T cells. Fluorescently labeled lipid-CpG (FAM) or biotinylated Pam_2_CSK_4_ (PE) was incubated for 1 hour either separately or together with splenic immune cells at 5 μM or 10 μg/mL, respectively. Splenic immune cells were gated by lineage markers to analyze depoting in **A)** B220^+^ B cells, **B)** CD4^+^ T cells, and **C)** CD8^+^ T cells. Loading was calculated as the fold increase of sample median fluorescent intensity (MFI) normalized to non-depoted cells for each ligand. p values as determined by one-way ANOVA and Tukey’s tests. *p < 0.05; ns, not significant. Data depict m ± s.d. (n = 3 independent samples).

Next, we optimized depoting conditions to more precisely control the abundance of ligands. Enzyme-linked immunosorbent assays (ELISAs) were used to quantify depoted lipid conjugates in purified mouse T cells. Conjugate concentration during depoting was tested over a 100-fold range and depoting times were tested ranging from 5 to 60 min. Cells were lysed after depoting to detect the average number of ligands per T cell. Depoting 5 μM of lipid-CpG for 60 min resulted in 25-fold more molecules per T cell when compared to depoting 0.05 μM of lipid-CpG (**Fig. 3A**).

We then quantified depoting of TLR2 ligands. Increasing depoting concentration of Pam_2_CSK_4_ from 0.1 to 10 μg/mL resulted in a 130-fold increase in the number of depoted molecules per T cell (**Fig. 3B, *left***). Both lipid-CpG and Pam_2_CSK_4_ depoting plateaued after 1 hour. Increasing depoting concentration did not change preferential insertion into CD4^+^ or CD8^+^ T cells (**Fig. S2A, S2B**). Pam_3_CSK_4_, a synthetic lipo-peptide with a triacyl palmitoyl lipid, is another well-defined TLR2 ligand (17, 29). We used this ligand to test if the third hydrophobic lipid tail altered depoting in T cells. Pam_3_CSK_4_ depoting increased with time, but increasing the depoting concentration of Pam_3_CSK_4_ from 0.1 μg/mL to 10 μg/mL resulted in a smaller 2.2-fold increase in number of depoted molecules per T cell (**Fig. 3B, *right***) when compared to Pam_2_CSK_4_. The diacyl and triacyl TLR2 ligands depoted into T-cell plasma membranes in a dose-dependent manner, suggesting that depoting is largely independent of tail number. We depoted cells in subsequent experiments for 1 hour at the highest ligand concentrations above to maximize depoting.

**Fig. 3:**
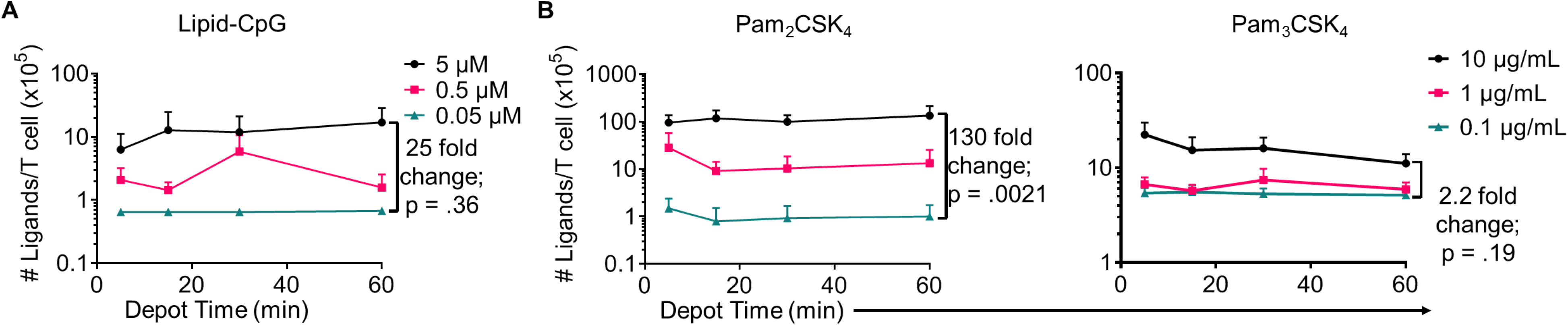
Depoted lipid-tailed TLR ligands into polyclonal T cells was dependent on input concentration for **A)** lipid-CpG; and **B)** Pam_2_CSK_4_ or Pam_3_CSK_4_ polypeptides. The fold increase in average ligand per T cell is indicated per ELISA analysis. Data depict mean ± s.d. (n = 3–4 independent samples).

### Depoted cell-surface ligands induced paracrine cell activation

TLR2 ligation has successfully enhanced cellular immunity by direct signaling on effector cells (e.g. T cells) and antigen-presenting cells (APCs)(17, 23). We hypothesized that TLR2 depoting could enhance T cell functions. To test this, we first validated that TLR2 ligands on T cells could engage its receptor on bystander cells using TLR2^+^ APCs. B cells were chosen as bystander APCs due to their dose-dependent sensitivity to Pam_2_CSK_4_ and Pam_3_CSK_4_ (Fig. S3A, S3B). T cells depoted with Pam_2_CSK_4_ activated B cells after 2 days of co-culture, upregulating MHC II (19-fold) and CD69 (3.8-fold) (**Fig. 4A**)(25). T cells depoted with Pam_3_CSK_4_ also activated B cells in co-culture, upregulating MHC II (21-fold) and CD69 (4.4-fold) (**Fig. 4B**). MHC II and CD69 expression levels activated by Pam_2_CSK_4_ and Pam_3_CSK_4_-depoted T cells were similar to 10 μg/mL of soluble ligand. This high dose of soluble ligand was 1000-5000-fold more concentrated than the estimated dose on depoted T cells (**Table S1**).

**Fig. 4:**
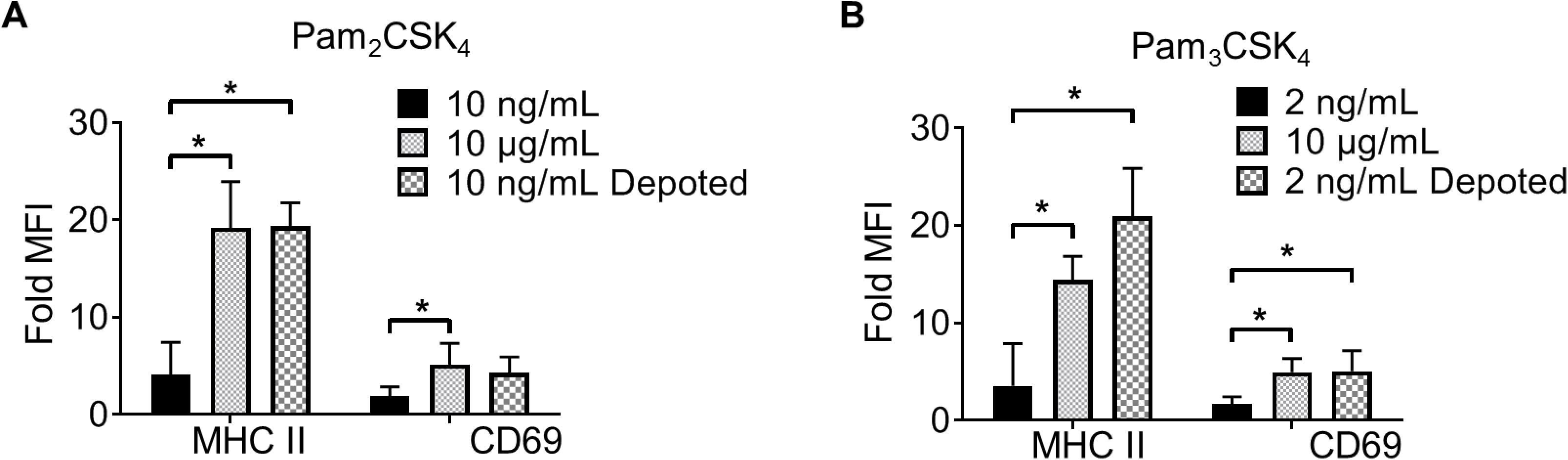
Depoted lipid-TLR ligands can activate cells through paracrine signaling. Polyclonal T cells were incubated with A) Pam_2_CSK_4_ or B) Pam_3_CSK_4_ for 1 h under indicated conditions, and then co-cultured with B cells for 2 days. B-cell activation was determined by fold MFI of activation markers, MHC II and CD69, measured by flow cytometry. Sample MFIs were then normalized to unstimulated B cell controls. n = 5–6 independent experiments, one-way ANOVA with Tukey’s tests; *p < 0.05; ns, not significant.

We next tested whether depoting can enhance cell sensitivity to TLR2 signaling per dose of ligand compared to soluble delivery. These doses (10 ng/mL for Pam_2_CSK_4_ and 2 ng/mL for Pam_3_CSK_4_) were matched to amount of respective ligand quantified in depoted cells as determined by ELISA (**Figs. 3B-C; Table S1**). T cells depoted with Pam_2_CSK_4_ induced higher expression of MHC II (5.3-fold) and CD69 (2.2-fold) when compared to the same dose of soluble Pam_2_CSK_4_ (**Fig. 4A**). Depoted Pam_3_CSK_4_ induced higher expression of MHC II (6.1-fold) and CD69 (2.8-fold) when compared to the same dose of soluble Pam_3_CSK_4_ (Fig. 4B). Consistent with our hypothesis, none of the dose-matched soluble Pam_2_CSK_4_ and Pam_3_CSK_4_ activated B cells. These studies demonstrated that depoted TLR2 ligands were presented to bystander cells, and that cell surface-bound presentation of TLR2 ligands provided stronger signaling than the equivalent ligand dose in solution.

### Intracellular ligands activated depoted immune cells but not bystanders

Bystander cells can be activated by paracrine interactions with depoted cells, as well as from release of depoted ligands over time into solution. We used lipid-CpG, a TLR9 ligand that must be internalized for signaling, to test whether release of depoted ligand over time contributed to paracrine cell activation. A co-culture assay was used with B cells as both depoted cells and bystander cells given their sensitivity to CpG stimulation (30). Lipid-CpG was depoted into B cells from wild-type mice, and then cocultured with congenic CD45.1^+^ bystander B cells for 2 days. Lipid-CpG depoted B cells showed increased expression of MHC II (19-fold) and CD69 (3.3-fold) when normalized to unstimulated B cells (**Fig. 5**; fold MFI). Levels of MHC II and CD69 were similar to those induced by a high dose of free CpG (5 μM) in solution. This confirmed that depoted ligand could be internalized to provide autocrine stimulation. We then analyzed activation of bystander cells to determine whether paracrine signaling occurred through soluble ligand release. Bystander CD45.1^+^ B cells were not activated, with activation marker levels comparable to unstimulated controls (**Fig. 5**). We analyzed whether ligand could spontaneously transfer from plasma membranes of depoted cells to undepoted bystanders. The presence of bystander B cells did not decrease activation of depoted B cells since MHC II and CD69 expression levels in cocultures were comparable to expression on depoted cells cultured alone. These data demonstrated that depoted ligand remained stably compartmentalized in depoted cells, and that lipid-CpG was not released at functional levels into solution or transferred to plasma membranes of bystander cells.

**Fig. 5:**
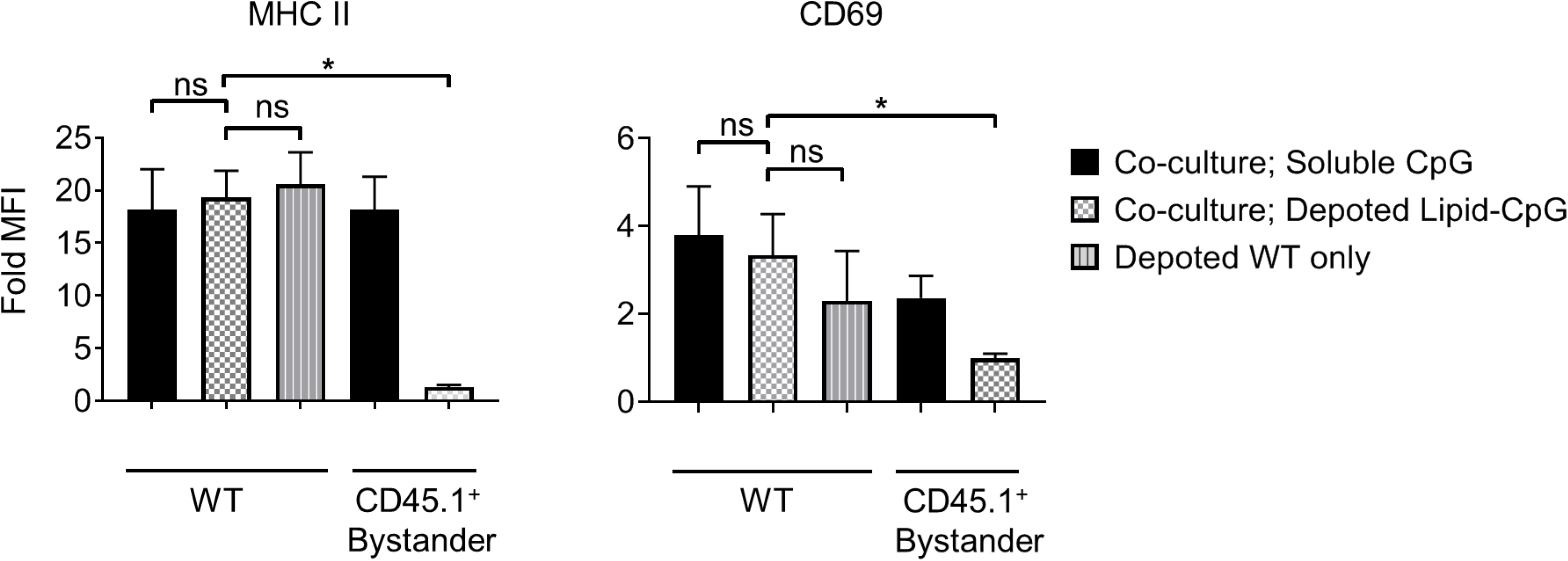
Depoted lipid-TLR ligands can activate cells through autocrine signaling. Wildtype (WT) B cells were depoted with 5 μM of lipid-CpG and co-cultured with CD45.1^+^ bystander B cells. MHCII and CD69 on both depoted (WT) and bystander (CD45.1^+^) B cells were measured after 2 days of co-culture by flow cytometry and normalized to respective unstimulated B cells (fold MFI). p values as determined by one-way ANOVA with Sidak’s multiple comparisons test; *p < 0.05; ns, not significant. Data depict m ± s.d. (n = 4 independent samples).

### Surface presentation of depoted TLR2 ligands was non-permanent

Depoting is a non-permanent cell modification, so we next characterized the persistence of depoted ligands on immune cell surfaces. Purified T cells were depoted with TLR2 ligands, and then rested for up to 8 days in IL-7. After resting, cells were fixed in 4% paraformaldehyde to preserve surface TLR2 ligands. The persistence of surface TLR2 ligands was determined by co-culturing fixed T cells with bystander B cells. Low level IL-7 supplementation was used to sustain T-cell viability without eliciting cell proliferation over the 8-day rest period (**Fig. S4**). We analyzed MHC II and CD69 levels on B cells induced by T cells fixed immediately after depoting (0-hour) and determined levels were comparable to levels observed in Fig. 3 (**Fig. 6A**). This verified that fixation did not alter the recognition of TLR2 ligand. Levels of surface-bound ligand decayed over time, with the shortest effective fast half-life of 0.49 days for Pam_3_CSK_4_ induction of CD69. Overall, bystander cell activation showed that surface-presented ligand functionally persisted between 2-4 days post-depoting.

Depoted ligands are not permanently persistent on cell surfaces, so their efficacy could be diluted as T cells divide. We used proliferating T cells to determine whether cell division causes faster decay of depoted ligands. Naïve T cells were primed for 2 days with concanavalin A and IL-7 before depoting. Primed cells were depoted with Pam_2_CSK_4_ or Pam_3_CSK_4_ and rested for up to 8 days. Primed T cells retained viability for 8 days in IL-7, similar to naïve T cells (**Fig. S4A, *right***). Depoted Pam_2_CSK_4_ and Pam_3_CSK_4_ decayed at a similar rate on primed T cells compared to naïve T cells, activating B cell bystanders as determined by MHC II and CD69 expression (**Fig. 6B**). The shortest effective fast half-life of 0.17 days was observed for MHC II levels stimulated by Pam3CSK4. Altogether, our data demonstrated that depoted TLR2 ligands persisted on plasma membranes of both naïve and primed T cells, providing functional paracrine signaling to bystander cells for multiple days. Proliferation of primed T cells did not further increase the surface decay rate for depoted ligands.

**Fig. 6:**
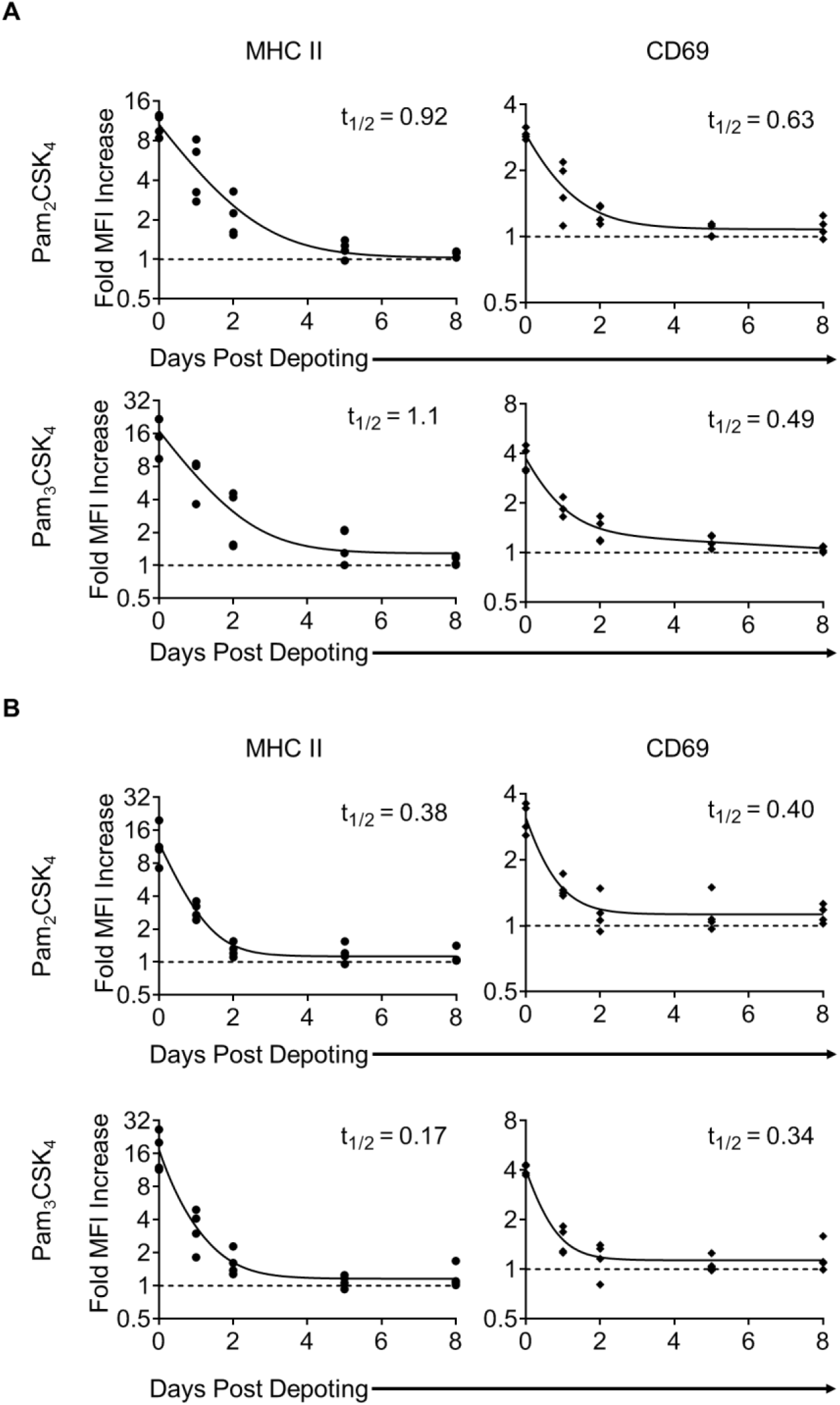
Surface presentation of depoted TLR2 ligands on T cells was non-permanent. Splenocytes were cultured either **A)** in absence of (naïve T cells), or **B)** presence of (primed T cells) 2 μg/mL of Concanavalin A and 2 ng/mL of IL-7 for 2 days. Purified T cells were then depoted with either Pam_2_CSK_4_ or Pam_3_CSK_4_ for 1 h at 37°C in supplemented RPMI media. T cells were then rested in IL-7 for 0, 1, 2, 5, or 8 days. At each timepoint, cells were fixed and co-cultured for 2 days with purified B cells. B-cell activation was determined by fold MFI of MHC II (circle) and CD69 (diamond). Solid lines are two-phase decay nonlinear regression curves as determined by independent replicate samples shown. Effective fast half-life in days (t_1/2_) as shown. Dashed lines represents baseline fold MFI of 1. n = 3–4 independent samples.

### Depoted lipid-conjugated TLR2 enhanced T cell activation

T cells do not constitutively express abundant levels of TLR2 or TLR9, but activated T cells rapidly increase TLR expression (31). Previous studies demonstrate TLR ligands can enhance T cell activation (17, 18, 32). Here, we tested whether depoting TLR ligands in T cells can enhance activation. We added TLR ligands in solution or depoted into pan CD4^+^ and CD8^+^ T cells from wild-type C57BL/6J mice. T cells were stimulated with αCD3/CD28-coated beads for 3 days, then analyzed for proliferation by flow cytometry. Depoting of TLR2 ligands induced more CD4^+^ T-cell proliferation compared to dose-matched soluble ligands (**Fig. 7A**). When T cells were depoted with Pam_3_CSK_4_ alone, CD4^+^ and CD8^+^ T-cell division indices were increased 1.9- and 2.9-fold, respectively, compared to soluble ligand (**Fig. 7B *top*, 7C top**). Depoting both TLR2 ligands increased CD4^+^ and CD8^+^ T-cell division indices by 2.2-fold and 3.3-fold, respectively, compared to soluble ligands. No lipid-conjugated ligands enhanced CD4^+^ T-cell proliferation index by depoting, but did enhance CD8^+^ T-cell proliferation index (**Fig. S5**). Altogether, these data demonstrated that Pam_3_CSK_4_, alone and in combination with other lipid-conjugated ligands, provided co-stimulation during T-cell division.

**Fig. 7:**
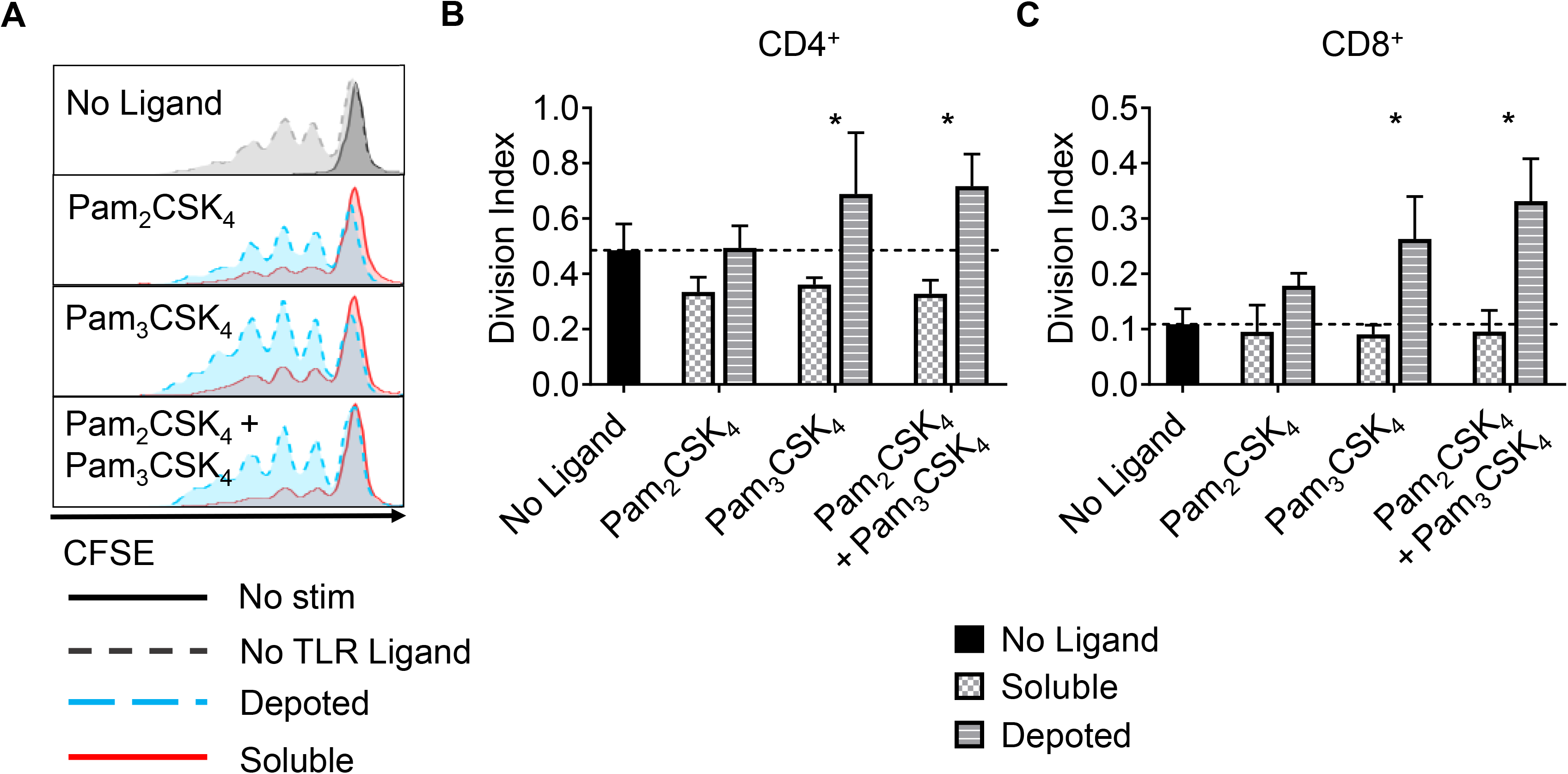
Depoted lipid-TLR2 ligand depoted enhanced proliferation of activated murine T cells. Purified polyclonal T cells were stained with 5 μM of carboxyfluorescein succinimidyl ester (CFSE). Different combinations of cell surface ligands (Pam_2_CSK_4_ and Pam_3_CSK_4_) were either directly added in solution (soluble) or depoted into polyclonal T cells for 1 h and cultured with αCD3/αCD28 beads for 3 days. **A)** Representative histograms of CD4^+^ T-cell proliferation from delivery of lipid-TLR2 ligand as measured by CFSE dilution. Quantification of division index of **B)** CD4+ and **C)** CD8+ T cells in bulk polyclonal T cells. Dotted lines represent respective averages (mean) of “No Ligand” conditions. p values between corresponding soluble vs depoted ligands as determined by one-tailed ratio paired t test; *p < 0.05; ns, not significant. Data depict m ± s.d. (n = 3 independent samples).

### Depoted TLR2 ligands enhanced T-cell activation by paracrine signaling

We tested whether depoting with TLR9 ligand could enhance T-cell activation and functions. Lipid-CpG did not increase proliferation of CD4^+^ or CD8^+^ T cells above levels compared to those without ligand. Combining lipid-CpG with Pam_2_CSK_4_ or Pam_3_CSK_4_ did not significantly enhance CD4^+^ T-cell proliferation, but did enhance CD8^+^ T-cell proliferation (**Fig. S6**). We determined how depoting with TLR2 and TLR9 ligands affected cytokine secretion by analyzing IL-2 and IL-4 levels as signature Th1 and Th2 cytokines, respectively. Soluble TLR9 ligand elicited detectable IL-4 production, while depoted TLR9 ligand decreased IL-4 levels below the limit of detection (**Fig. S7A**). Soluble and depoted TLR2 ligands resulted in IL-4 levels below the limit of detection. Depoting of TLR2 and TLR9 ligands resulted in higher IL-2 levels compared to dose-matched soluble ligands (**Fig. S7B**). These data suggested that co-stimulation of T cells by depoted TLR ligands elicited a more pro-inflammatory Th1-like phenotype.

While depoting increased T-cell proliferation, we next analyzed whether depoted TLR2 ligands provide autocrine co-stimulation or paracrine co-stimulation to bystander T cells. Depoted T cells were mixed with non-depoted bystander T cells, and co-cultures were stimulated with αCD3/CD28 beads for 3 days. CD4^+^ and CD8^+^ T-cell division indices of Pam_3_CSK_4_-depoted cells increased 1.6-fold compared to respective soluble ligand (**Fig. 8A**). Cis-depoted Pam_2_CSK_4_ and Pam_3_CSK_4_ (depoted onto the same cell) increased division indices of depoted cells: 1.8-fold for CD4^+^ T cells and 1.7-fold for CD8^+^ T cells compared to soluble ligands. No changes were observed in proliferation indices (**Fig. S8**). Pam_3_CSK_4_-depoted T cells also increased bystander CD4^+^ and CD8^+^ T-cell division indices by 1.3-fold compared to soluble ligand. Cis-depoted Pam_2_CSK_4_ and Pam_3_CSK_4_ increased division indices of bystander cells by 1.4-fold for CD4^+^ T cells and 1.3-fold for CD8^+^ T cells compared to soluble ligands. These data demonstrated that TLR2 ligand depoted T cells could engage in autocrine and paracrine enhancement of proliferation, activating themselves as well as bystander T cells. We analyzed whether depoted T cells could enhance cytokine secretion by paracrine engagement of bystander T cells. No differences in secretion of Th1 cytokines, IFNγ or IL-2, were observed with addition of bystander T cells to depoted T cells (**Fig. 8B**). These data showed that depoted TLR2 ligands enhanced bystander T-cell proliferation but not Th1 cytokine secretion.

**Fig. 8:**
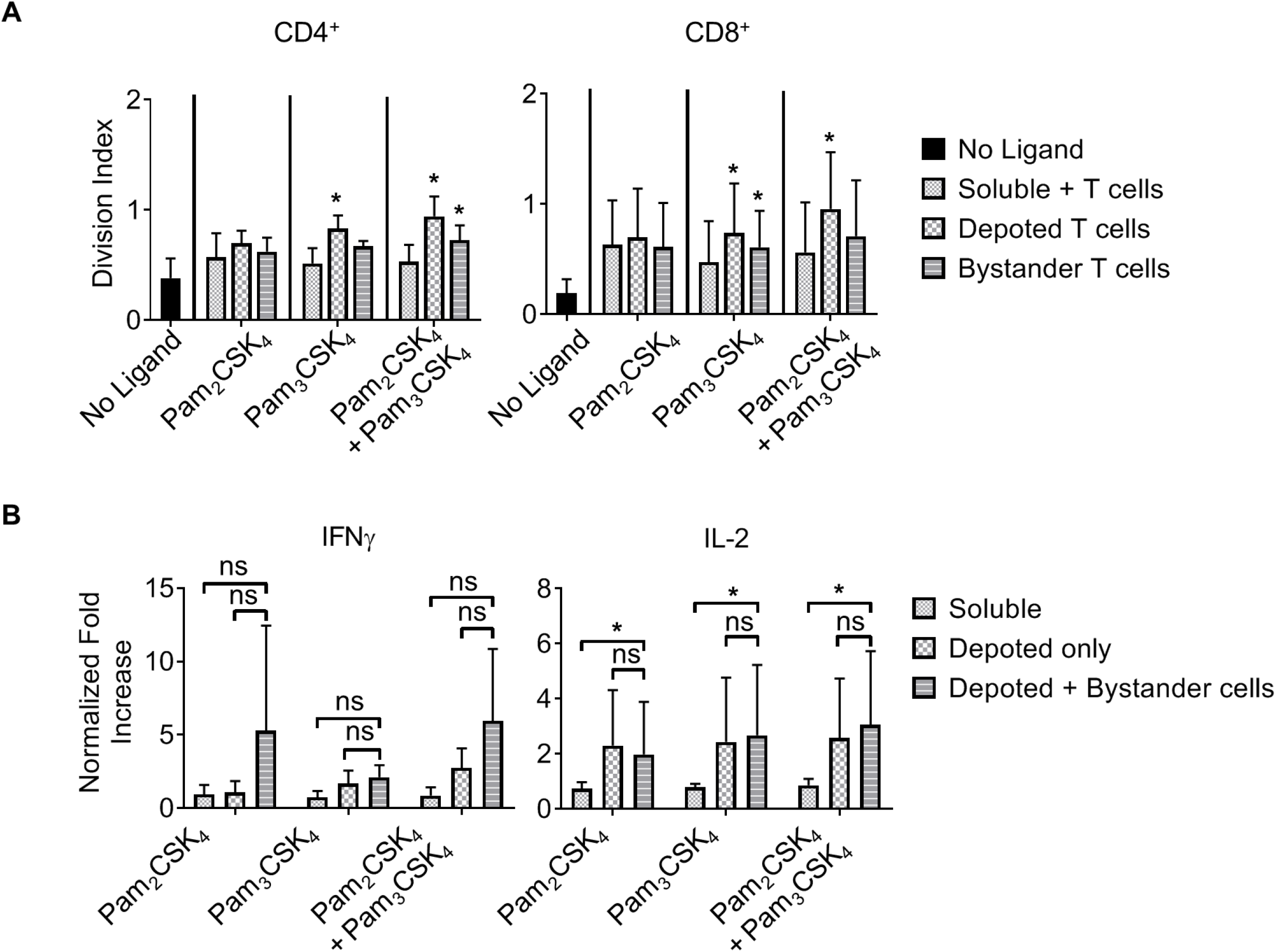
Depoted lipid-TLR2 ligands enhanced murine T-cell activation. **A)** Purified polyclonal T cells were stained with 5 μM of carboxyfluorescein succinimidyl ester (CFSE). Different combinations of cell surface ligands (Pam_2_CSK_4_ and Pam_3_CSK_4_) were either directly added in solution (soluble) or depoted into polyclonal T cells, and co-cultured with non-depoted T cells and αCD3/αCD28 beads for 3 days. Quantification of division index of CD4^+^ and CD8^+^ T cells in bulk polyclonal T cells. p values within each treatment group is determined by comparing with respective soluble ligand as determined by one-way ANOVA with Holm-Sidak’s multiple comparisons tests; *p < 0.05; ns, not significant. Data depict m ± s.d. (n = 5 independent samples). **B)** IFNγ and IL-2 from cell supernatents were measured by ELISA on day 2. Concentration values were normalized to respective cytokine concentration of αCD3/αCD28 bead-stimulated T cells in the absence of TLR2 ligand. p values within each treatment group as determined by two-way ANOVA with Sidak’s multiple comparisons test; *p < 0.05; ns, not significant. Data depict m ± s.d. (n = 4 independent samples for IFNγ; n = 5 independent samples for IL-2).

However, depoting of TLR2 ligands increased IFNγ and IL-2 secretion levels when compared to dose-matched soluble ligands. Depoting TLR2 ligands increased, but not significantly, IFNγ levels (**Fig. 8B, *left***). Depoting TLR2 ligands increased IL-2 levels and the IL-2 receptor, CD25, expression (**Fig. 8B, right; Fig. S9**). Depoting with Pam_2_CSK_4_ induced a 2.7-fold increase IL-2 levels compared to soluble ligand (**Fig. 8B**). Depoting with Pam_3_CSK_4_ induced a 3.4-fold increase in IL-2 levels compared to soluble ligand. *Cis*-depoted Pam_2_CSK_4_ and Pam_3_CSK_4_ induced a 3.4-fold increase in IL-2 levelscompared to soluble ligands. These data again demonstrated that surface-bound presentation of depoted TLR2 ligands provided stronger signaling than dose-matched soluble ligands through enhanced IL-2 secretion.

## Discussion

In this study, we characterized depoting of lipid conjugates as a novel method for simple, non-genetic engineering of immune cells. Our data showed diacyl C16 and C18, and triacyl C16 lipid tails conjugated to DNA and protein cargoes depoted into plasma membranes in a concentration-dependent manner. We used TLR2 and TLR9 ligands to demonstrate the potential to engineer cells with ligands targeting surface and intracellular receptors, respectively. Depoted TLR2 ligands enhanced bystander B- and T-cell activation when compared to dose-matched soluble ligands. Depoted TLR9 ligands showed autocrine activation of depoted cells with minimal paracrine activation of bystanders, demonstrating delivery of depoted ligands with minimal release into solution or transfer between plasma membranes. Depoting stored detectable levels of surface-bound ligands on resting and proliferating T cells for bystander cell activation for similar amounts of time, up to 4 days of *in vitro* co-culture. We also demonstrated multiplex loading of 2 distinct ligands onto the same plasma membranes without reaching loading saturation or decreasing cell viability. Altogether, our results demonstrated lipid-mediated depoting is a facile, modular drug delivery platform for diverse ligands with receptors in distinct subcellular spaces.

Depoting TLR ligands can provide precise control over ligand dose by changing input ligand concentration. Varying the depoting concentration of lipid-CpG and Pam2CSK4 over a 100-fold range resulted in 25-fold and 130-fold increases, respectively, of ligand on cells. This suggests that changing depoting concentrations could modulate dose control, a feature that can dictate whether a drug induces immunogenic or tolerogenic responses (33, 34). We determined that cells could be loaded by multiplexing with 2 depoted ligands and have no detectable loading saturation. However, differential loading of B and T cells in **Fig. 2** suggest that lymphocyte subsets have distinct plasma membrane compositions, including TLR9 ligand-internalization receptors that may skew immune cell surface binding and alter total depoting capacity in a ligand- and cell-dependent manner (35, 36). Here, we demonstrated depoting of >1 million molecules per cell for some ligands, which is classified as very abundant based on recent estimates (37). This shows that depoting loads a high absolute amount of cargoes onto plasma membranes beyond the likely capacity of just surface receptor binding.

We also demonstrated that the lipids we used (i.e., diacyl/triacyl C16 and diacyl C18 tails) effectively delivered ligands to plasma membranes. Our previous study showed that a single acyl lipid tail is inefficient at TLR ligand delivery (8, 11). Here, our results demonstrated that two and three-tailed lipids depoted efficiently. Thus, multiple lipid tails are a key design parameter for membrane delivery, and may be a simple way to enhance delivery of cargo to plasma membranes. One explanation of lipid-mediated insertion of TLR2 lipid-peptides, i.e. Pam_2_CSK_4_ and Pam_3_CSK_4_, into plasma membranes is their spontaneous formation of micelles when delivered at a high ligand concentration. This may result in particulate-mediated fusion with plasma membranes. However, the critical micelle concentrations of TLR2 ligands are much higher than our input concentrations of 10 μg/mL (38). Our hypothesis of lipid-ligand “depoting” is a likely explanation that is supported by lipid-mediated partitioning of plasma membranes (27, 28) and the natural membrane insertion of GPI-anchored proteins. Altogether, depoting is advantageous not only for delivering synthetic TLR2 and TLR9 ligands as shown in our study, but may also apply to other immunostimulatory TLR ligands with greater than three lipid tails, including TLR4 ligands.

The controlled delivery of autocrine or paracrine signaling is critical to the further understanding of immunological signaling and enhancing cell-based therapies. We demonstrated that depoting of a cell surface ligand enhanced CD4+ and CD8+ T cell division, and also promoted inflammatory responses of bystander T cells. This simple method of delivery can be leveraged in the clinic, including activating TLR2 in cytotoxic cells (23, 39). In cancer, prolonging the persistence of patient-infused T cells while also eliciting inflammatory responses by reprogramming endogenous tumor-infiltrating lymphocytes or anti-tumor macrophages can enhance therapeutic efficacy. However, paracrine signaling of delivered adjuvants is not always desirable for promoting anti-tumor responses (40, 41). Our delivery platform also demonstrated exclusive autocrine signaling with an intracellular ligand by targeting cell endosomes while avoiding bystander activation. This feature is critical for minimizing off-target effects by rational selection of clinically relevant intracellular ligands paired with target cell types. Controlled autocrine signaling may also help dissect underlying mechanisms of poorly understood disease pathogenesis including graft-versus-host disease (GVHD). In hematopoietic stem cell transplantation, the role of TLR ligation in donor vs recipient cells is unclear in GVHD despite clinical protocols that remove TLR-activating microbes (42, 43). Limiting TLR signaling to exclusively autocrine by depoting donor cells may illuminate donor vs graft responses in a more controlled manner.

The use of genetically modified cells for cell-based immunotherapies has been one of the most promising, fastest-developing therapeutic approaches in recent years. However, there is a critical need to improve efficacy while reducing toxicity (44, 45). In 2017, the FDA approved 2 adoptive T-cell immunotherapies using chimeric antigen receptors (CARs) for treating cancer, with hundreds of other T-cell therapies currently in clinical trials in cancer and other chronic diseases (46). The addition of more co-stimulatory signals, other adjuvants, or signaling domains to CARs by genetic additions has become an increasing trend in the field (47–49). However, the uncontrolled release of genetically encoded inflammatory cytokines or other adjuvants from perpetually activated CAR T cells presents a new challenge for toxicity management: lack of dose control (45, 50). New CAR technologies will likely entail even more genetic additions, which invariably will carry genetically-related adverse responses and toxicities. These genetic encodings prolong this adverse risks over the lifetime of the cell (51, 52). Expanding genetic modifications to future therapeutic genes is also limited by the packaging capacity of viral vectors (53). We propose that depoting can address these challenges by complementing genetic engineering by providing non-permanent delivery of co-stimulatory signals for 2–4 days. While depoted cargoes are inherently transient, this time window is critical for therapeutic efficacy. For example, CAR T cells can initially kill a large proportion of target cells within 2-3 days post-infusion (54, 55). Further, Pam_3_CSK_4_ can reactivate latently infected HIV reservoirs in patients to enhance recognition and eradication of infected cells (56). TLR2 surface presentation by depoted T cells may enhance this “shock and kill” response by localizing ligand activity and enhancing ligand potency of cytotoxic T cells to reactivate the latent reservoir. Rational selection and combination of depoted vs. genetically engineered features can improve CAR T-cell persistence and efficacy *in vivo* while minimizing toxicities that may constrain clinical efficacy.

Drug delivery strategies for cellular engineering, from gene editing to nanoparticle and lipid carriers, are under continuous and rapid development (2, 57, 58). For example, nanoparticles have been used in multiple formulations to boost cellular therapies by delivering drugs with immune cells *in vivo* (59–61). Previous studies have shown palmitoyl lipid-tailed conjugates of CD28 ligand or IL-2 cytokine can be “painted” onto T cell surfaces to enhance cell function (9, 10). Our study significantly expands upon these results by demonstrating the modularity of lipid-mediated delivery for different tail lengths and numbers, and showing that delivery of lipid-ligands can be achieved to cell surface and intracellular receptors. A current limitation of lipid-mediated depoting is the transient nature of the ligand on the cell surface, which will limit duration of ligand signaling on therapeutic cells against chronic diseases such as cancer. Future studies should focus on determining design criteria to effectively depot other ligand combinations with longer persistence or higher abundance to optimize immunogenic or immunosuppressive effects *in vitro* and *in vivo* in disease models.

Our data prove that depoting can engineer diverse immune cell types and expands existing knowledge on lipid dynamics that can be used for designing future drug delivery approaches. We used two innate TLRs targeted to directly enhance T-cell immunotherapies, and showed enhanced T-cell responses with depoted lipid-TLR ligands. This enables re-examination of pathways for T-cell activation that have been previously understudied due to delivery barriers. Rational selection of more clinically potent ligands for autocrine or paracrine signaling will illustrate the utility of depoting in a broad array of cell-based therapies.

## Acknowledgements

Flow cytometry analyses were performed at the University of Maryland School of Medicine Center for Innovative Biomedical Resources, Flow Cytometry Shared Service – Baltimore, Maryland. Confocal analyses were performed at the Keith Porter Imaging Facility. We thank Yun Jiao, Drs. Ryan Pearson and Jennie Leach for critical reading and feedback on the manuscript. This work was supported by a grant from the Elsa U Pardee Foundation (to GLS), UMBC’s Undergraduate Research Awards (to EMS and GS), UMBC Summer Faculty Fellowship (to GLS), a Supplement for Undergraduate Research Experiences (to GLS and EMS), and the Commercialization & ENTR REsearch (CENTRE) Funding Initiative of the Alex Brown Center for Entrepreneurship at UMBC (to GLS). GLS is funded in part by the UMGCC P30 grant under award number P30 CA134274 from the National Cancer Institute, NIH. EMS was supported in part by the Nathan Schnaper Intern Program in Translational Cancer Research (NIH R25CA186872).

## References

1. Barton GM, Kagan JC, Medzhitov R. Intracellular localization of Toll-like receptor 9 prevents recognition of self DNA but facilitates access to viral DNA. Nat Immunol. 2006;7(1):49–56. Epub 2005/12/13. doi: 10.1038/ni1280. PubMed PMID: 16341217.

2. Stewart MP, Langer R, Jensen KF. Intracellular Delivery by Membrane Disruption: Mechanisms, Strategies, and Concepts. Chem Rev. 2018;118(16):7409–531. Epub 2018/07/28. doi: 10.1021/acs.chemrev.7b00678. PubMed PMID: 30052023; PubMed Central PMCID: PMCPMC6763210.

3. Kunath K, von Harpe A, Fischer D, Petersen H, Bickel U, Voigt K, et al. Low-molecular-weight polyethylenimine as a non-viral vector for DNA delivery: comparison of physicochemical properties, transfection efficiency and in vivo distribution with high-molecular-weight polyethylenimine. J Control Release. 2003;89(1):113–25. Epub 2003/04/16. doi: 10.1016/s0168-3659(03)00076-2. PubMed PMID: 12695067.

4. Olden BR, Cheng Y, Yu JL, Pun SH. Cationic polymers for non-viral gene delivery to human T cells. J Control Release. 2018;282:140–7. Epub 2018/03/09. doi: 10.1016/j.jconrel.2018.02.043. PubMed PMID: 29518467; PubMed Central PMCID: PMCPMC6008197.

5. Nayerossadat N, Maedeh T, Ali PA. Viral and nonviral delivery systems for gene delivery. Adv Biomed Res. 2012;1:27. Epub 2012/12/05. doi: 10.4103/2277-9175.98152. PubMed PMID: 23210086; PubMed Central PMCID: PMCPMC3507026.

6. Smith TT, Stephan SB, Moffett HF, McKnight LE, Ji W, Reiman D, et al. In situ programming of leukaemia-specific T cells using synthetic DNA nanocarriers. Nat Nanotechnol. 2017;12(8):813–20. Epub 2017/04/19. doi: 10.1038/nnano.2017.57. PubMed PMID: 28416815; PubMed Central PMCID: PMCPMC5646367.

7. Park K. Facing the truth about nanotechnology in drug delivery. ACS Nano. 2013;7(9):7442–7. Epub 2014/02/05. doi: 10.1021/nn404501g. PubMed PMID: 24490875; PubMed Central PMCID: PMCPMC3914010.

8. Liu H, Moynihan KD, Zheng Y, Szeto GL, Li AV, Huang B, et al. Structure-based programming of lymph-node targeting in molecular vaccines. Nature. 2014;507(7493):519–22. Epub 2014/02/18. doi: 10.1038/nature12978. PubMed PMID: 24531764; PubMed Central PMCID: PMCPMC4069155.

9. Zheng G, Liu S, Wang P, Xu Y, Chen A. Arming tumor-reactive T cells with costimulator B7-1 enhances therapeutic efficacy of the T cells. Cancer Res. 2006;66(13):6793–9. Epub 2006/07/05. doi: 10.1158/0008-5472.can-06-0435. PubMed PMID: 16818656.

10. Chou SH, Shetty AV, Geng Y, Xu L, Munirathinam G, Pipathsouk A, et al. Palmitate-derivatized human IL-2: a potential anticancer immunotherapeutic of low systemic toxicity. Cancer Immunol Immunother. 2013;62(3):597–603. Epub 2012/11/06. doi: 10.1007/s00262-012-1364-8. PubMed PMID: 23124508; PubMed Central PMCID: PMCPMC4393711.

11. Liu H, Kwong B, Irvine DJ. Membrane anchored immunostimulatory oligonucleotides for in vivo cell modification and localized immunotherapy. Angew Chem Int Ed Engl.2011;50(31):7052–5. Epub 2011/06/21. doi: 10.1002/anie.201101266. PubMed PMID:21688362; PubMed Central PMCID: PMCPMC3166645.

12. Yu C, An M, Li M, Liu H. Immunostimulatory Properties of Lipid Modified CpG Oligonucleotides. Mol Pharm. 2017;14(8):2815–23. Epub 2017/07/08. doi: 10.1021/acs.molpharmaceut.7b00335. PubMed PMID: 28686452.

13. Adams S. Toll-like receptor agonists in cancer therapy. Immunotherapy. 2009;1(6):949–64. Epub 2010/06/22. doi: 10.2217/imt.09.70. PubMed PMID: 20563267; PubMed Central PMCID: PMCPMC2886992.

14. Zhang Y, Luo F, Cai Y, Liu N, Wang L, Xu D, et al. TLR1/TLR2 agonist induces tumor regression by reciprocal modulation of effector and regulatory T cells. J Immunol. 2011;186(4):1963–9. Epub 2011/01/11. doi: 10.4049/jimmunol.1002320. PubMed PMID: 21217015.

15. Uehori J, Matsumoto M, Tsuji S, Akazawa T, Takeuchi O, Akira S, et al. Simultaneous blocking of human Toll-like receptors 2 and 4 suppresses myeloid dendritic cell activation induced by Mycobacterium bovis bacillus Calmette-Guerin peptidoglycan. Infect Immun. 2003;71(8):4238–49. Epub 2003/07/23. doi: 10.1128/iai.71.8.4238-4249.2003. PubMed PMID: 12874299; PubMed Central PMCID: PMCPMC165983.

16. Lamm DL, Blumenstein BA, Crawford ED, Montie JE, Scardino P, Grossman HB, et al. A randomized trial of intravesical doxorubicin and immunotherapy with bacille Calmette-Guerin for transitional-cell carcinoma of the bladder. N Engl J Med. 1991;325(17):1205–9. Epub 1991/11/03. doi: 10.1056/nejm199110243251703. PubMed PMID: 1922207.

17. Geng D, Zheng L, Srivastava R, Velasco-Gonzalez C, Riker A, Markovic SN, et al. Amplifying TLR-MyD88 signals within tumor-specific T cells enhances antitumor activity to suboptimal levels of weakly immunogenic tumor antigens. Cancer Res. 2010;70(19):7442–54. Epub 2010/09/03. doi: 10.1158/0008-5472.can-10-0247. PubMed PMID: 20807806; PubMed Central PMCID: PMCPMC3463001.

18. Wong RM, Smith KA, Tam VL, Pagarigan RR, Meisenburg BL, Quach AM, et al. TLR-9 signaling and TCR stimulation co-regulate CD8(+) T cell-associated PD-1 expression. Immunol Lett. 2009;127(1):60–7. Epub 2009/09/16. doi: 10.1016/j.imlet.2009.09.002. PubMed PMID: 19751765.

19. Nierkens S, den Brok MH, Roelofsen T, Wagenaars JA, Figdor CG, Ruers TJ, et al. Route of administration of the TLR9 agonist CpG critically determines the efficacy of cancer immunotherapy in mice. PLoS One. 2009;4(12):e8368. Epub 2009/12/19. doi: 10.1371/journal.pone.0008368. PubMed PMID: 20020049; PubMed Central PMCID: PMCPMC2791230.

20. Bendigs S, Salzer U, Lipford GB, Wagner H, Heeg K. CpG-oligodeoxynucleotides co-stimulate primary T cells in the absence of antigen-presenting cells. Eur J Immunol. 1999;29(4):1209–18. Epub 1999/05/06. doi: 10.1002/(sici)1521-4141(199904)29:04<1209::aid-immu1209>3.0.co;2-j. PubMed PMID: 10229088.

21. Davila E, Velez MG, Heppelmann CJ, Celis E. Creating space: an antigen-independent, CpG-induced peripheral expansion of naive and memory T lymphocytes in a full T-cell compartment. Blood. 2002;100(7):2537–45. Epub 2002/09/20. doi: 10.1182/blood-2002-02-0401. PubMed PMID: 12239167.

22. Ribas A, Medina T, Kummar S, Amin A, Kalbasi A, Drabick JJ, et al. SD-101 in Combination with Pembrolizumab in Advanced Melanoma: Results of a Phase Ib, Multicenter Study. Cancer Discov. 2018;8(10):1250–7. Epub 2018/08/30. doi: 10.1158/2159-8290.cd-18-0280. PubMed PMID: 30154193; PubMed Central PMCID: PMCPMC6719557.

23. Lu H, Yang Y, Gad E, Inatsuka C, Wenner CA, Disis ML, et al. TLR2 agonist PSK activates human NK cells and enhances the antitumor effect of HER2-targeted monoclonal antibody therapy. Clin Cancer Res. 2011;17(21):6742–53. Epub 2011/09/16. doi: 10.1158/1078-0432.ccr-11-1142. PubMed PMID: 21918170; PubMed Central PMCID: PMCPMC3206987.

24. Thomas M, Ponce-Aix S, Navarro A, Riera-Knorrenschild J, Schmidt M, Wiegert E, et al. Immunotherapeutic maintenance treatment with toll-like receptor 9 agonist lefitolimod in patients with extensive-stage small-cell lung cancer: results from the exploratory, controlled, randomized, international phase II IMPULSE study. Ann Oncol. 2018;29(10):2076–84. Epub 2018/08/24. doi: 10.1093/annonc/mdy326. PubMed PMID: 30137193; PubMed Central PMCID: PMCPMC6225892.

25. Szeto GL, Van Egeren D, Worku H, Sharei A, Alejandro B, Park C, et al. Microfluidic squeezing for intracellular antigen loading in polyclonal B-cells as cellular vaccines. Sci Rep. 2015;5:10276. Epub 2015/05/23. doi: 10.1038/srep10276. PubMed PMID: 25999171; PubMed Central PMCID: PMCPMC4441198.

26. Avalos AM, Ploegh HL. Early BCR Events and Antigen Capture, Processing, and Loading on MHC Class II on B Cells. Front Immunol. 2014;5:92. Epub 2014/03/22. doi: 10.3389/fimmu.2014.00092. PubMed PMID: 24653721; PubMed Central PMCID: PMCPMC3948085.

27. Ong S, Liu H, Qiu X, Bhat G, Pidgeon C. Membrane partition coefficients chromatographically measured using immobilized artificial membrane surfaces. Anal Chem. 1995;67(4):755–62. Epub 1995/02/15. doi: 10.1021/ac00100a011. PubMed PMID: 7702190.

28. Jacobs RE, White SH. The nature of the hydrophobic binding of small peptides at the bilayer interface: implications for the insertion of transbilayer helices. Biochemistry. 1989;28(8):3421–37. Epub 1989/04/18. doi: 10.1021/bi00434a042. PubMed PMID: 2742845.

29. Krishnegowda G, Hajjar AM, Zhu J, Douglass EJ, Uematsu S, Akira S, et al. Induction of proinflammatory responses in macrophages by the glycosylphosphatidylinositols of Plasmodium falciparum: cell signaling receptors, glycosylphosphatidylinositol (GPI) structural requirement, and regulation of GPI activity. J Biol Chem. 2005;280(9):8606–16. Epub 2004/12/30. doi: 10.1074/jbc.M413541200. PubMed PMID: 15623512; PubMed Central PMCID: PMCPMC4984258.

30. Hacein-Bey-Abina S, Garrigue A, Wang GP, Soulier J, Lim A, Morillon E, et al. Insertional oncogenesis in 4 patients after retrovirus-mediated gene therapy of SCID-X1. J Clin Invest. 2008;118(9):3132–42. Epub 2008/08/09. doi: 10.1172/jci35700. PubMed PMID: 18688285; PubMed Central PMCID: PMCPMC2496963.

31. Hornung V, Rothenfusser S, Britsch S, Krug A, Jahrsdorfer B, Giese T, et al. Quantitative expression of toll-like receptor 1-10 mRNA in cellular subsets of human peripheral blood mononuclear cells and sensitivity to CpG oligodeoxynucleotides. J Immunol. 2002;168(9):4531–7. Epub 2002/04/24. doi: 10.4049/jimmunol.168.9.4531. PubMed PMID: 11970999.

32. Komai-Koma M, Jones L, Ogg GS, Xu D, Liew FY. TLR2 is expressed on activated T cells as a costimulatory receptor. Proc Natl Acad Sci U S A. 2004;101(9):3029–34. Epub 2004/02/26. doi: 10.1073/pnas.0400171101. PubMed PMID: 14981245; PubMed Central PMCID: PMCPMC365739.

33. Ray A, Chakraborty K, Ray P. Immunosuppressive MDSCs induced by TLR signaling during infection and role in resolution of inflammation. Front Cell Infect Microbiol. 2013;3:52. Epub 2013/09/26. doi: 10.3389/fcimb.2013.00052. PubMed PMID: 24066282; PubMed Central PMCID: PMCPMC3776133.

34. Volpi C, Fallarino F, Pallotta MT, Bianchi R, Vacca C, Belladonna ML, et al. High doses of CpG oligodeoxynucleotides stimulate a tolerogenic TLR9-TRIF pathway. Nat Commun. 2013;4:1852. Epub 2013/05/16. doi: 10.1038/ncomms2874. PubMed PMID: 23673637.

35. Lahoud MH, Ahmet F, Zhang JG, Meuter S, Policheni AN, Kitsoulis S, et al. DEC-205 is a cell surface receptor for CpG oligonucleotides. Proc Natl Acad Sci U S A. 2012;109(40):16270–5. Epub 2012/09/19. doi: 10.1073/pnas.1208796109. PubMed PMID: 22988114; PubMed Central PMCID: PMCPMC3479608.

36. Hacker H, Mischak H, Miethke T, Liptay S, Schmid R, Sparwasser T, et al. CpG-DNA-specific activation of antigen-presenting cells requires stress kinase activity and is preceded by non-specific endocytosis and endosomal maturation. Embo j. 1998;17(21):6230–40. Epub 1998/11/03. doi: 10.1093/emboj/17.21.6230. PubMed PMID: 9799232; PubMed Central PMCID: PMCPMC1170949.

37. Wisniewski JR, Hein MY, Cox J, Mann M. A “proteomic ruler” for protein copy number and concentration estimation without spike-in standards. Mol Cell Proteomics. 2014;13(12):3497–506. Epub 2014/09/17. doi: 10.1074/mcp.M113.037309. PubMed PMID: 25225357; PubMed Central PMCID: PMCPMC4256500.

38. Hamley IW, Kirkham S, Dehsorkhi A, Castelletto V, Reza M, Ruokolainen J. Toll-like receptor agonist lipopeptides self-assemble into distinct nanostructures. Chem Commun (Camb). 2014;50(100):15948–51. Epub 2014/11/11. doi: 10.1039/c4cc07511k. PubMed PMID: 25382300.

39. Lu H, Yang Y, Gad E, Wenner CA, Chang A, Larson ER, et al. Polysaccharide krestin is a novel TLR2 agonist that mediates inhibition of tumor growth via stimulation of CD8 T cells and NK cells. Clin Cancer Res. 2011;17(1):67–76. Epub 2010/11/12. doi: 10.1158/1078-0432.ccr-10-1763. PubMed PMID: 21068144; PubMed Central PMCID: PMCPMC3017241.

40. Akkaya M, Akkaya B, Kim AS, Miozzo P, Sohn H, Pena M, et al. Toll-like receptor 9 antagonizes antibody affinity maturation. Nat Immunol. 2018;19(3):255–66. Epub 2018/02/25. doi: 10.1038/s41590-018-0052-z. PubMed PMID: 29476183; PubMed Central PMCID: PMCPMC5839995.

41. Moseman EA, Liang X, Dawson AJ, Panoskaltsis-Mortari A, Krieg AM, Liu YJ, et al. Human plasmacytoid dendritic cells activated by CpG oligodeoxynucleotides induce the generation of CD4+CD25+ regulatory T cells. J Immunol. 2004;173(7):4433–42. Epub 2004/09/24. doi: 10.4049/jimmunol.173.7.4433. PubMed PMID: 15383574.

42. Heidegger S, van den Brink MR, Haas T, Poeck H. The role of pattern-recognition receptors in graft-versus-host disease and graft-versus-leukemia after allogeneic stem cell transplantation. Front Immunol. 2014;5:337. Epub 2014/08/08. doi: 10.3389/fimmu.2014.00337. PubMed PMID: 25101080; PubMed Central PMCID: PMCPMC4102927.

43. Penack O, Holler E, van den Brink MR. Graft-versus-host disease: regulation by microbe-associated molecules and innate immune receptors. Blood. 2010;115(10):1865–72. Epub 2010/01/01. doi: 10.1182/blood-2009-09-242784. PubMed PMID: 20042727.

44. Rosenberg SA, Restifo NP. Adoptive cell transfer as personalized immunotherapy for human cancer. Science. 2015;348(6230):62–8. Epub 2015/04/04. doi: 10.1126/science.aaa4967. PubMed PMID: 25838374; PubMed Central PMCID: PMCPMC6295668.

45. Bonifant CL, Jackson HJ, Brentjens RJ, Curran KJ. Toxicity and management in CAR T-cell therapy. Mol Ther Oncolytics. 2016;3:16011. Epub 2016/09/15. doi: 10.1038/mto.2016.11. PubMed PMID: 27626062; PubMed Central PMCID: PMCPMC5008265.

46. Home - ClinicalTrials.gov 2019. Available from: https://clinicaltrials.gov/.

47. Rafiq S, Yeku OO, Jackson HJ, Purdon TJ, van Leeuwen DG, Drakes DJ, et al. Targeted delivery of a PD-1-blocking scFv by CAR-T cells enhances anti-tumor efficacy in vivo. Nat Biotechnol. 2018;36(9):847–56. Epub 2018/08/14. doi: 10.1038/nbt.4195. PubMed PMID: 30102295; PubMed Central PMCID: PMCPMC6126939.

48. Zhao Z, Condomines M, van der Stegen SJC, Perna F, Kloss CC, Gunset G, et al. Structural Design of Engineered Costimulation Determines Tumor Rejection Kinetics and Persistence of CAR T Cells. Cancer Cell. 2015;28(4):415–28. Epub 2015/10/16. doi: 10.1016/j.ccell.2015.09.004. PubMed PMID: 26461090; PubMed Central PMCID: PMCPMC5003056.

49. Schubert ML, Hoffmann JM, Dreger P, Muller-Tidow C, Schmitt M. Chimeric antigen receptor transduced T cells: Tuning up for the next generation. Int J Cancer. 2018;142(9):1738–47. Epub 2017/11/10. doi: 10.1002/ijc.31147. PubMed PMID: 29119551.

50. Neelapu SS, Tummala S, Kebriaei P, Wierda W, Locke FL, Lin Y, et al. Toxicity management after chimeric antigen receptor T cell therapy: one size does not fit ‘ALL’. Nat Rev Clin Oncol. 2018;15(4):218. Epub 2018/02/13. doi: 10.1038/nrclinonc.2018.20. PubMed PMID: 29434334; PubMed Central PMCID: PMCPMC6716606.

51. Essand M, Loskog AS. Genetically engineered T cells for the treatment of cancer. J Intern Med. 2013;273(2):166–81. Epub 2012/12/04. doi: 10.1111/joim.12020. PubMed PMID: 23198862; PubMed Central PMCID: PMCPMC3607417.

52. Ruella M, Xu J, Barrett DM, Fraietta JA, Reich TJ, Ambrose DE, et al. Induction of resistance to chimeric antigen receptor T cell therapy by transduction of a single leukemic B cell. Nat Med. 2018;24(10):1499–503. Epub 2018/10/03. doi: 10.1038/s41591-018-0201-9. PubMed PMID: 30275568; PubMed Central PMCID: PMCPMC6511988.

53. Wang X, Riviere I. Clinical manufacturing of CAR T cells: foundation of a promising therapy. Mol Ther Oncolytics. 2016;3:16015. Epub 2016/06/28. doi: 10.1038/mto.2016.15. PubMed PMID: 27347557; PubMed Central PMCID: PMCPMC4909095.

54. Herzig E, Kim KC, Packard TA, Vardi N, Schwarzer R, Gramatica A, et al. Attacking Latent HIV with convertibleCAR-T Cells, a Highly Adaptable Killing Platform. Cell. 2019; 179(4):880–94.e10. Epub 2019/11/02. doi: 10.1016/j.cell.2019.10.002. PubMed PMID: 31668804.

55. Anurathapan U, Chan RC, Hindi HF, Mucharla R, Bajgain P, Hayes BC, et al. Kinetics of tumor destruction by chimeric antigen receptor-modified T cells. Mol Ther. 2014;22(3):623–33. Epub 2013/11/12. doi: 10.1038/mt.2013.262. PubMed PMID: 24213558; PubMed Central PMCID: PMCPMC3945803.

56. Novis CL, Archin NM, Buzon MJ, Verdin E, Round JL, Lichterfeld M, et al. Reactivation of latent HIV-1 in central memory CD4(+) T cells through TLR-1/2 stimulation. Retrovirology. 2013;10:119. Epub 2013/10/26. doi: 10.1186/1742-4690-10-119. PubMed PMID: 24156240; PubMed Central PMCID: PMCPMC3826617.

57. Csizmar CM, Petersburg JR, Wagner CR. Programming Cell-Cell Interactions through Non-genetic Membrane Engineering. Cell Chem Biol. 2018;25(8):931–40. Epub 2018/06/19. doi: 10.1016/j.chembiol.2018.05.009. PubMed PMID: 29909993; PubMed Central PMCID: PMCPMC6470397.

58. Neelapu SS, Tummala S, Kebriaei P, Wierda W, Gutierrez C, Locke FL, et al. Chimeric antigen receptor T-cell therapy - assessment and management of toxicities. Nat Rev Clin Oncol. 2018;15(1):47–62. Epub 2017/09/20. doi: 10.1038/nrclinonc.2017.148. PubMed PMID: 28925994; PubMed Central PMCID: PMCPMC6733403.

59. Bourquin C, Anz D, Zwiorek K, Lanz AL, Fuchs S, Weigel S, et al. Targeting CpG oligonucleotides to the lymph node by nanoparticles elicits efficient antitumoral immunity. J Immunol. 2008;181(5):2990–8. Epub 2008/08/21. doi: 10.4049/jimmunol.181.5.2990. PubMed PMID: 18713969.

60. McHugh MD, Park J, Uhrich R, Gao W, Horwitz DA, Fahmy TM. Paracrine co-delivery of TGF-beta and IL-2 using CD4-targeted nanoparticles for induction and maintenance of regulatory T cells. Biomaterials. 2015;59:172–81. Epub 2015/05/15. doi: 10.1016/j.biomaterials.2015.04.003. PubMed PMID: 25974747; PubMed Central PMCID: PMCPMC5997248.

61. Kwong B, Liu H, Irvine DJ. Induction of potent anti-tumor responses while eliminating systemic side effects via liposome-anchored combinatorial immunotherapy. Biomaterials. 2011;32(22):5134–47. Epub 2011/04/26. doi: 10.1016/j.biomaterials.2011.03.067. PubMed PMID: 21514665; PubMed Central PMCID: PMCPMC3140866.

