## Supplemental Figures for "Lipid-mediated insertion of Toll-like receptor (TLR) ligands for facile immune cell engineering"

**Fig. S1**

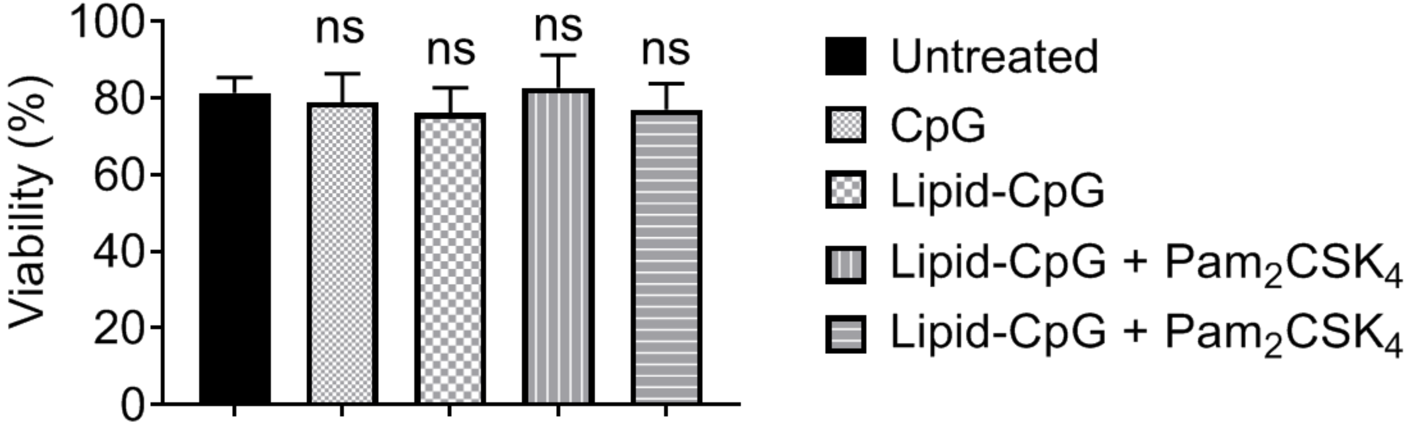

100
80
60
40
20
0
Viability (%)
ns
ns
ns
ns
Untreated
CpG
Lipid-CpG
Lipid-CpG + Pam2CSK4
Lipid-CpG + Pam2CSK4

Fig. S2

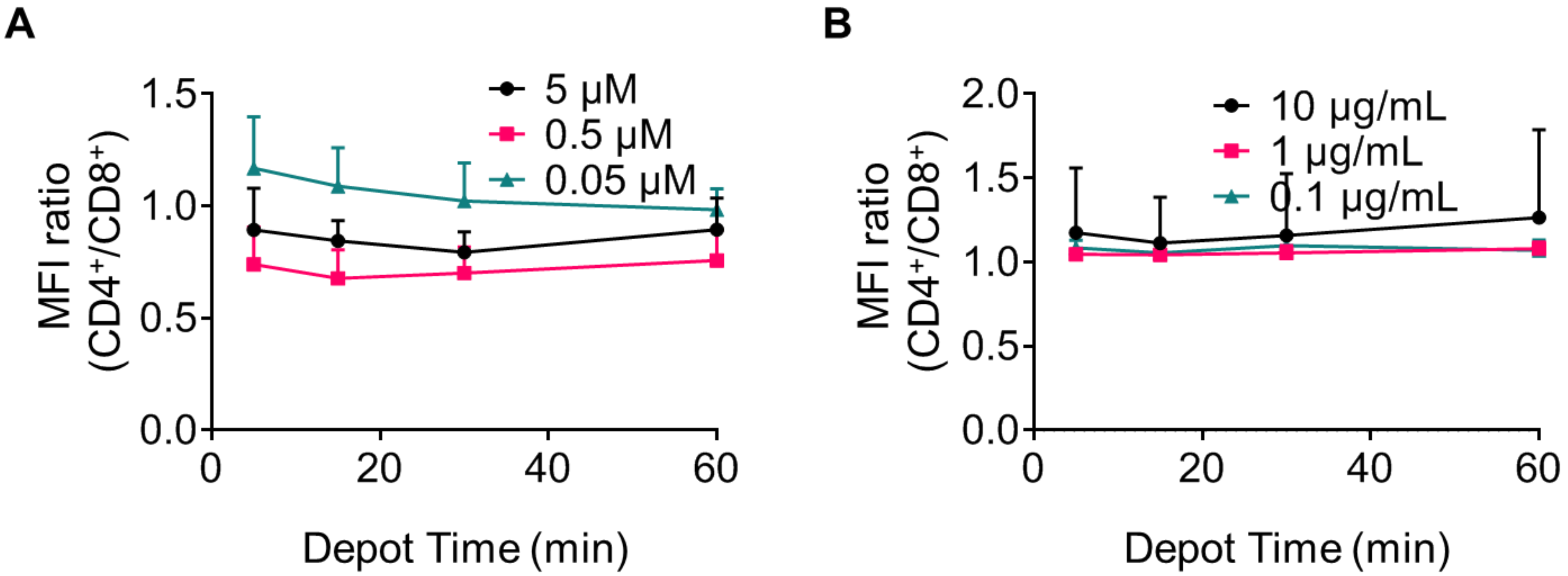
A
B
MFI ratio (CD4+/CD8+)
Depot Time (min)
5 μM
0.5 μM
0.05 μM
10 μg/mL
1 μg/mL
0.1 μg/mL
1.5
1.0
0.5
0.0
2.0
0
20
40
60

### Table S1

| TLR2 Ligand | # Ligands/T cell | Molecular Weight (g/mol) | ng/T cell | ng/50,000 T cells | ng/mL |
|---|---|---|---|---|---|
| $Pam_2CSK_4$ | 1.E+07 | 1724.44 | 4.E-05 | 2 | 10 |
| $Pam_3CSK_4$ | 2.E+06 | 1962.9 | 7.E-06 | 0.4 | 2 |

**Fig. S3**

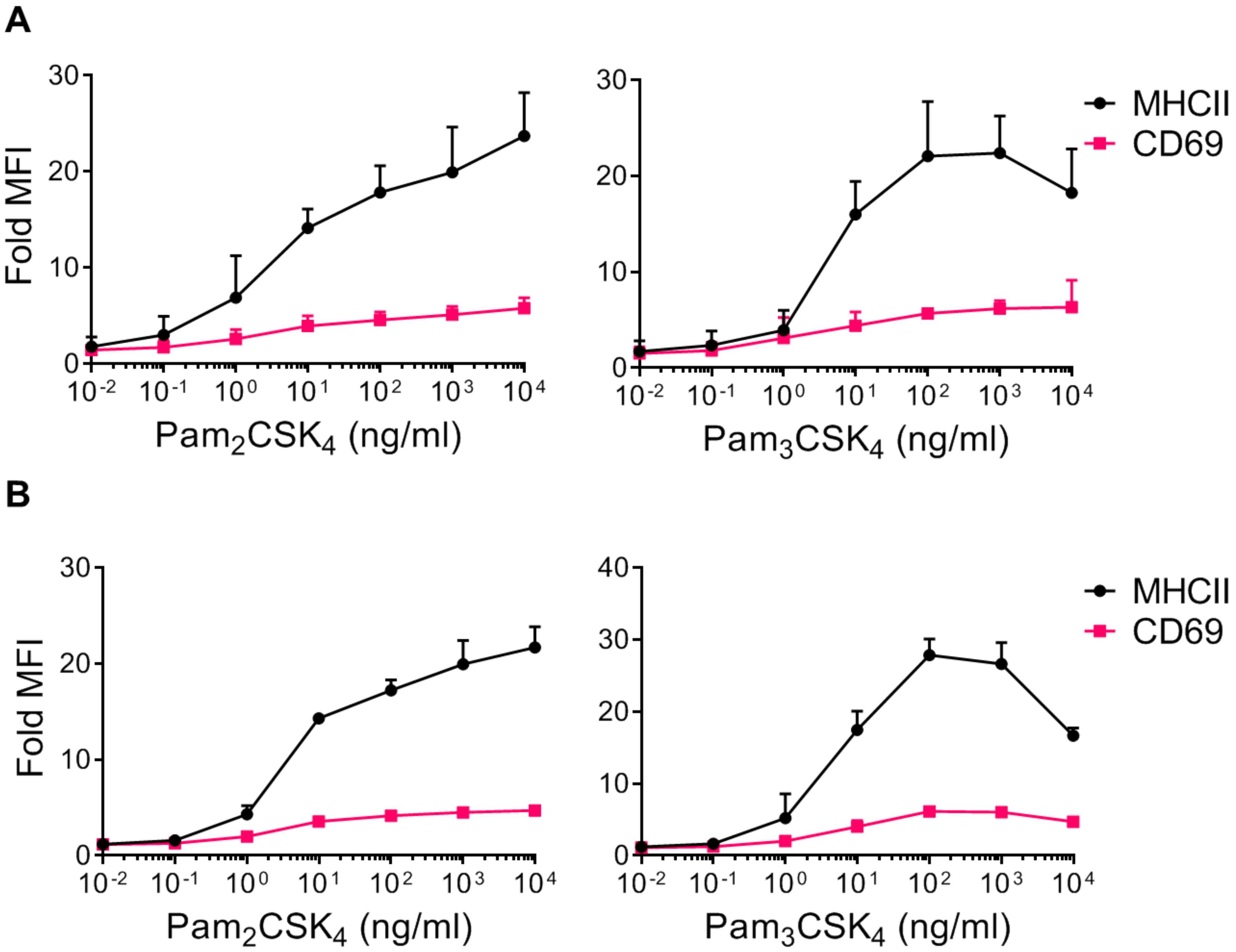

A
Fold MFI
30
20
10
0
10-2 10-1 100 101 102 103 104
Pam2CSK4 (ng/ml)
Pam3CSK4 (ng/ml)
MHCII
CD69
B
Fold MFI
40
Pam2CSK4 (ng/ml)
Pam3CSK4 (ng/ml)
MHCII
CD69

Fig. S4

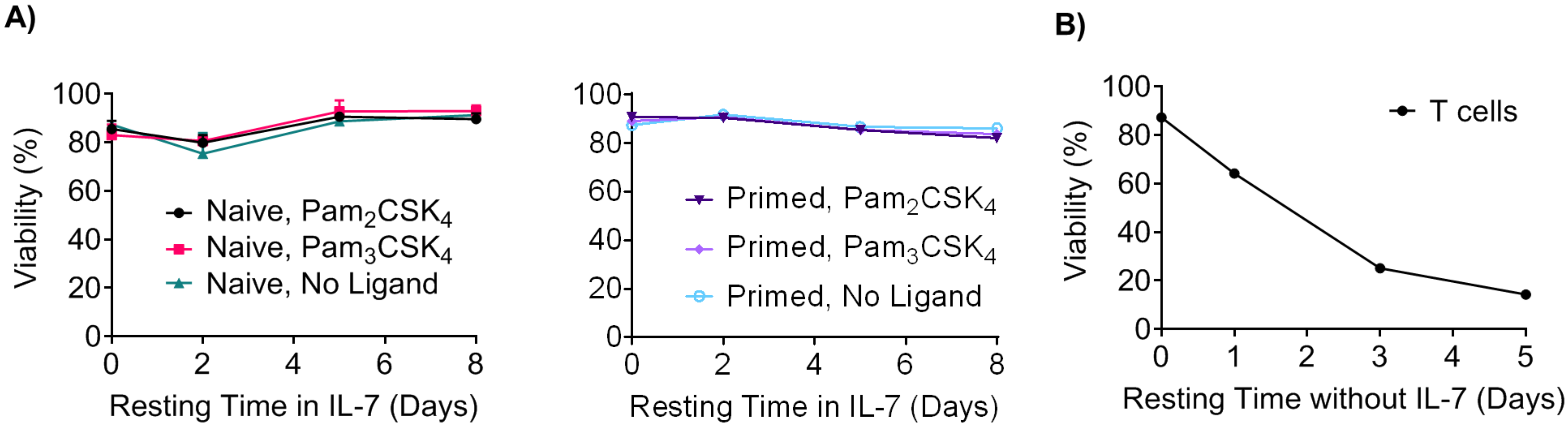
A)
B)
Viability (%)
0
20
40
60
80
100
Naive, $Pam_2CSK_4$
Naive, $Pam_3CSK_4$
Naive, No Ligand
0
2
4
6
8
Resting Time in IL-7 (Days)
Primed, $Pam_2CSK_4$
Primed, $Pam_3CSK_4$
Primed, No Ligand
Resting Time in IL-7 (Days)
T cells
0
1
2
3
4
5
Resting Time without IL-7 (Days)

**Fig. S5**

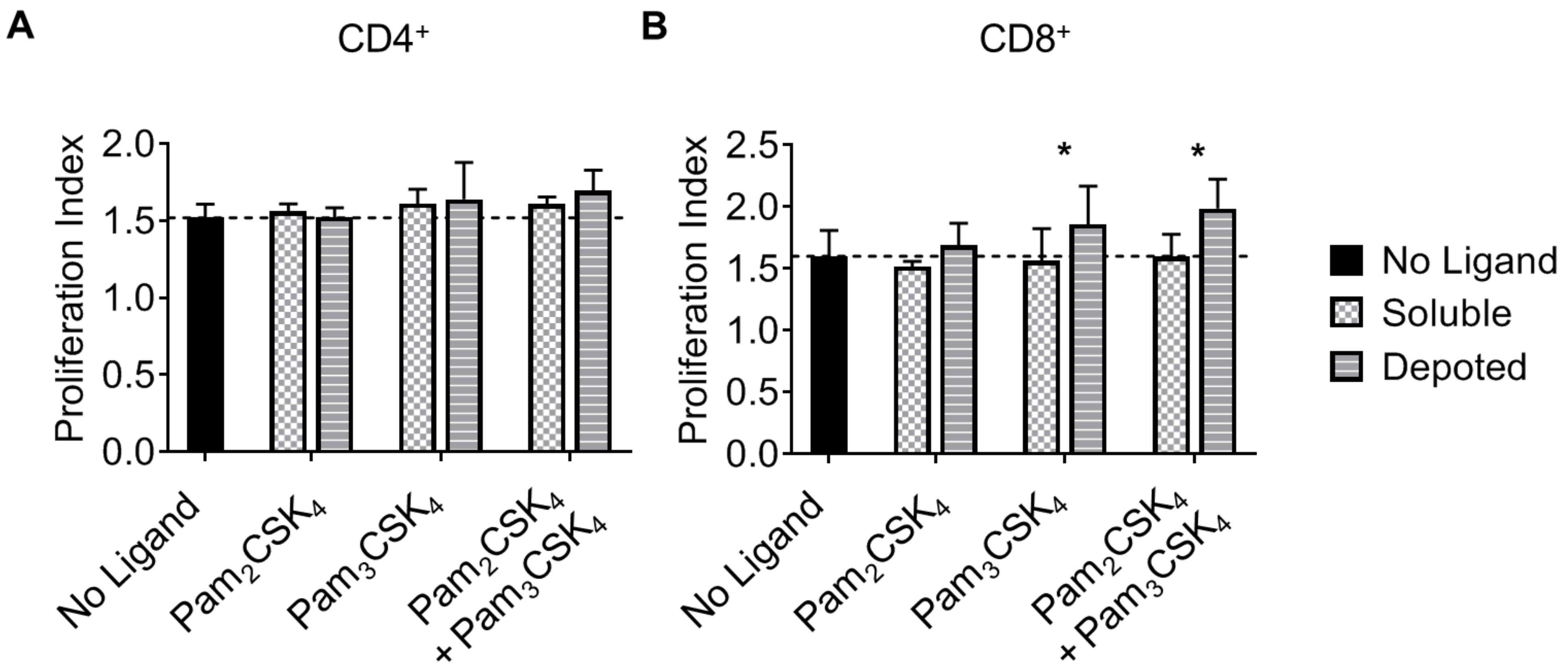
A
CD4+
Proliferation Index
2.0
1.5
1.0
0.5
0.0
No Ligand
Pam2CSK4
Pam3CSK4
Pam2CSK4
+ Pam3CSK4
B
CD8+
Proliferation Index
2.5
2.0
1.5
1.0
0.5
0.0
*
*
No Ligand
Pam2CSK4
Pam3CSK4
Pam2CSK4
+ Pam3CSK4
No Ligand
Soluble
Depoted

### Fig. S6

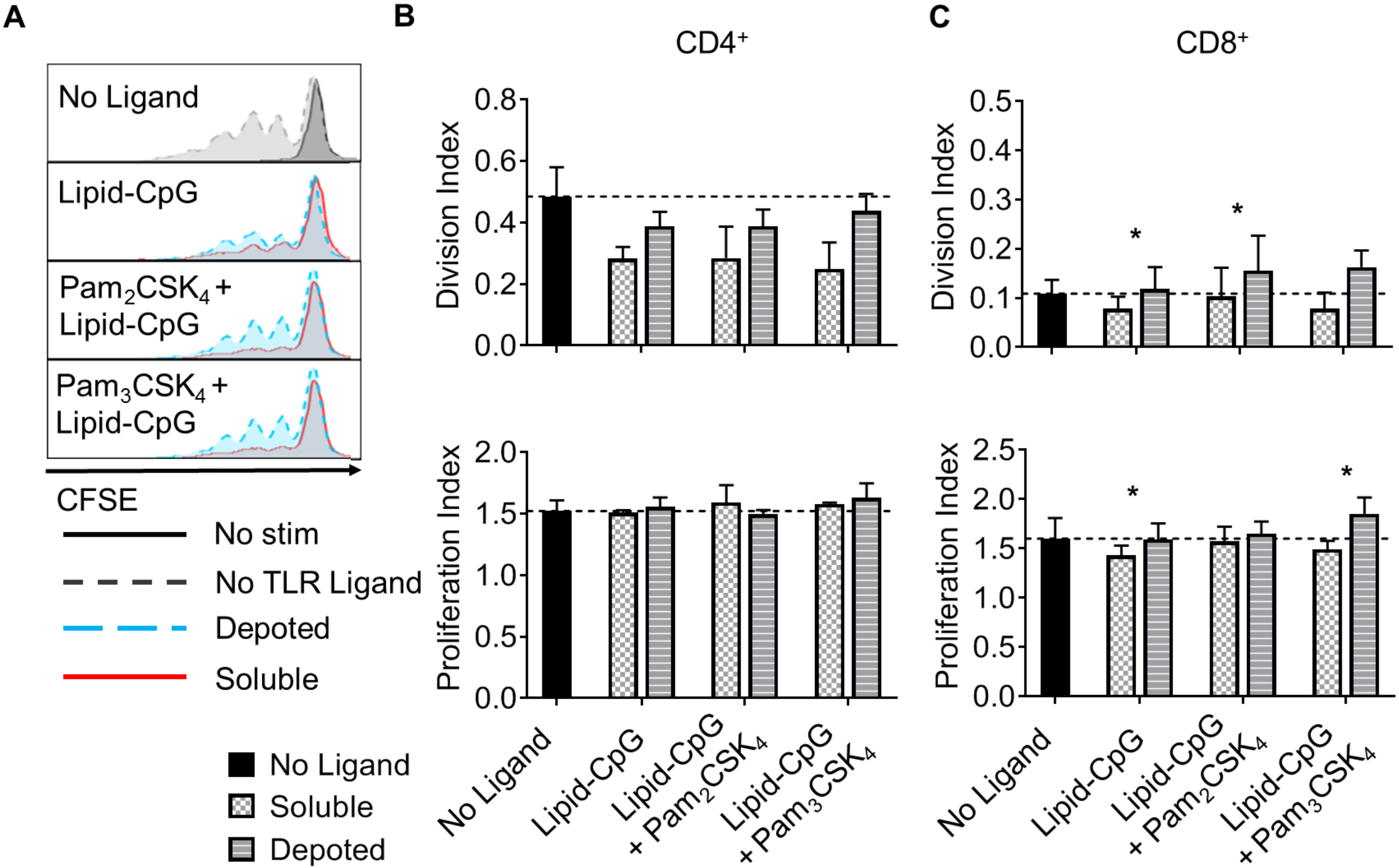
A
No Ligand
Lipid-CpG
$Pam_2CSK_4$ +
Lipid-CpG
$Pam_3CSK_4$ +
Lipid-CpG
CFSE
No stim
No TLR Ligand
Depoted
Soluble
B
$CD4^+$
Division Index
0.0
0.2
0.4
0.6
0.8
Proliferation Index
0.0
0.5
1.0
1.5
2.0
C
$CD8^+$
Division Index
0.0
0.1
0.2
0.3
0.4
0.5
*
*
Proliferation Index
0.0
0.5
1.0
1.5
2.0
2.5
*
*
No Ligand
Lipid-CpG
Lipid-CpG + $Pam_2CSK_4$
Lipid-CpG + $Pam_3CSK_4$
No Ligand
Soluble
Depoted

**Fig. S7**

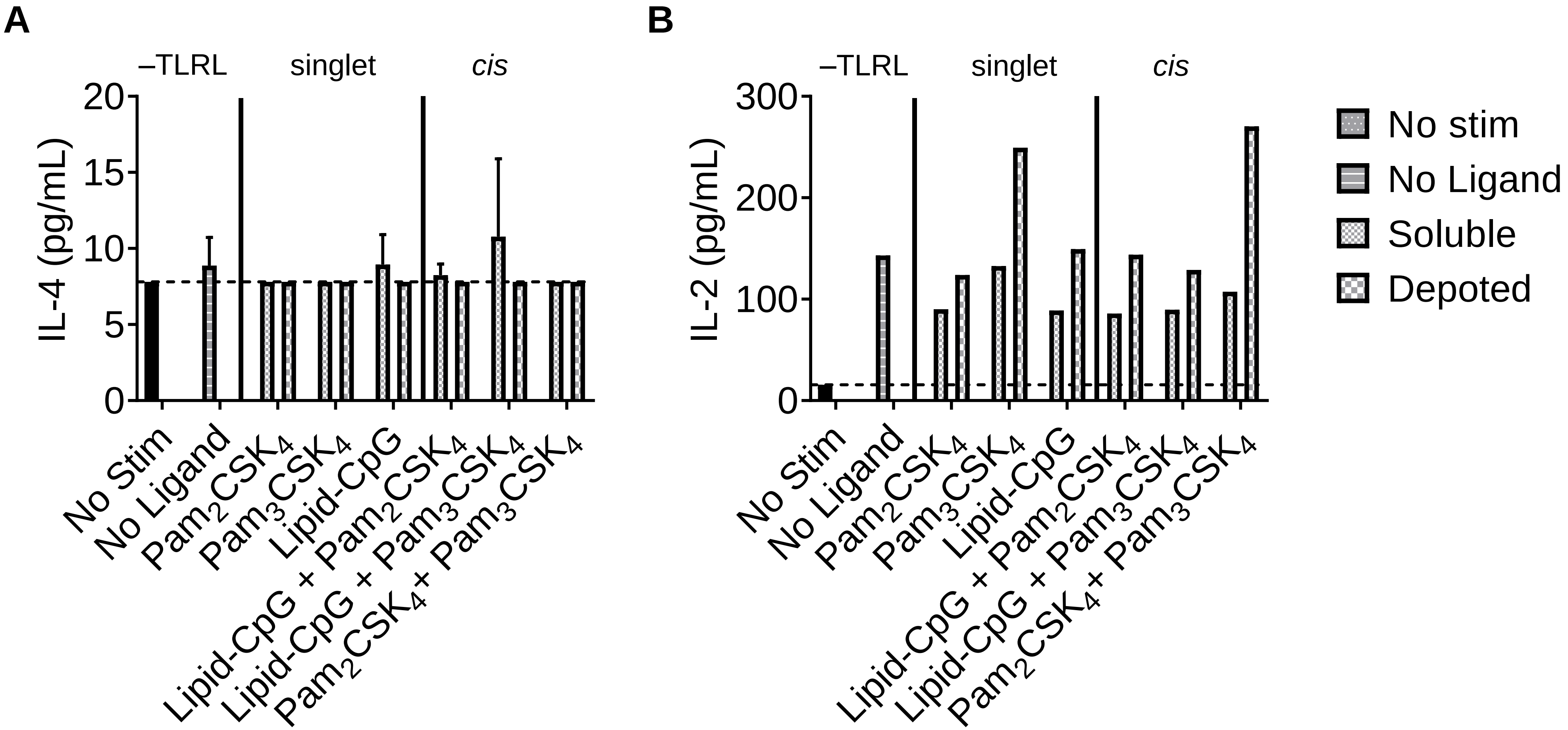
A
–TLRL
singlet
cis
IL-4 (pg/mL)
20
15
10
5
0
No Stim
No Ligand
Pam2CSK4
Pam3CSK4
Lipid-CpG
Lipid-CpG + Pam2CSK4
Lipid-CpG + Pam3CSK4
Pam2CSK4+ Pam3CSK4
B
–TLRL
singlet
cis
IL-2 (pg/mL)
300
200
100
0
No Stim
No Ligand
Pam2CSK4
Pam3CSK4
Lipid-CpG
Lipid-CpG + Pam2CSK4
Lipid-CpG + Pam3CSK4
Pam2CSK4+ Pam3CSK4
No stim
No Ligand
Soluble
Depoted

**Fig. S8**

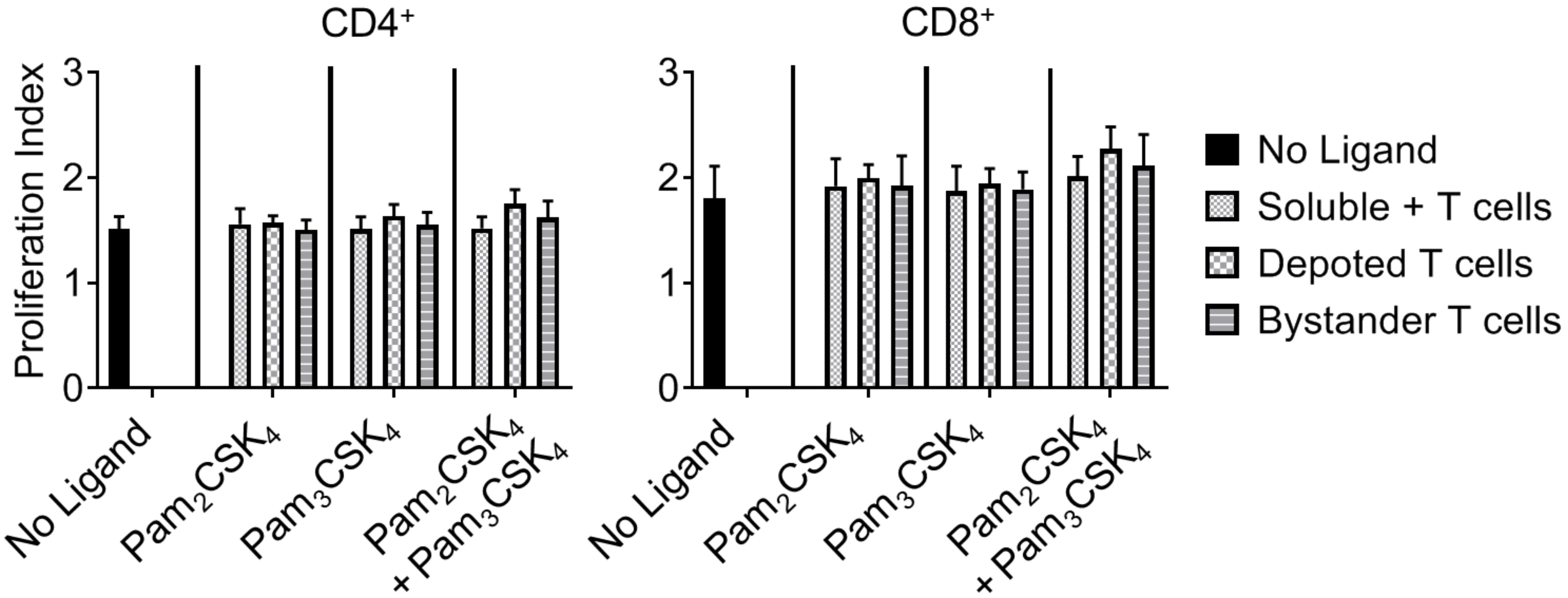
CD4⁺
CD8⁺
Proliferation Index
0
1
2
3
No Ligand
Pam2CSK4
Pam3CSK4
Pam2CSK4 + Pam3CSK4
No Ligand
Soluble + T cells
Depoted T cells
Bystander T cells

**Fig. S9**

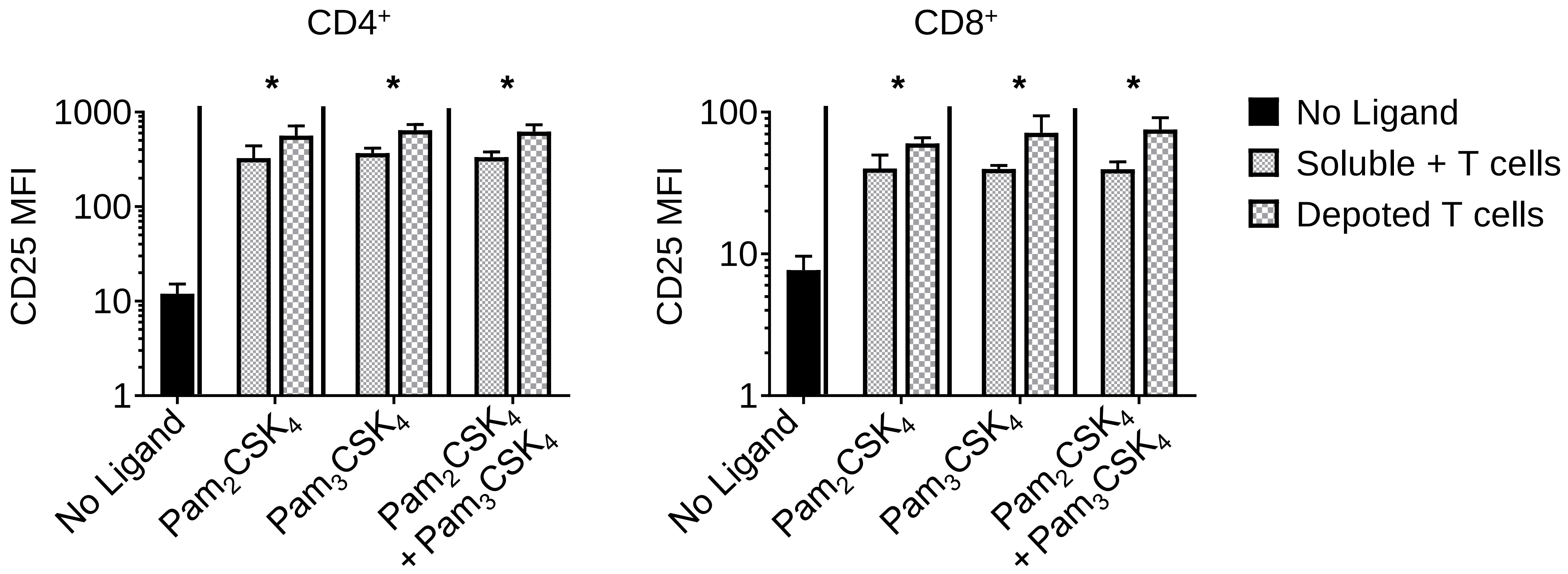
CD4+
CD8+
*
*
*
*
*
*
1000
100
10
1
CD25 MFI
100
10
1
CD25 MFI
No Ligand
Pam2CSK4
Pam3CSK4
Pam2CSK4
+ Pam3CSK4
No Ligand
Pam2CSK4
Pam3CSK4
Pam2CSK4
+ Pam3CSK4
No Ligand
Soluble + T cells
Depoted T cells
