## Supplemental Figure Legends for "Lipid-mediated insertion of Toll-like receptor (TLR) ligands for facile immune cell engineering"

**Fig. S1**: Viability of murine immune cells after TLR ligand binding. Cell viability of each treatment group as measured by DAPI staining. Indicated p values as determined by one-way ANOVA and Dunnett’s test when compared to “Untreated”; *p < 0.05; ns, not significant. Data depict m ± s.d. (n = 3 independent samples).

**Fig. S2**: Depoted lipid-tailed TLR ligands in is unchanged in ratio of CD4^+^ to CD8^+^ T cells for **A)** lipid-CpG, and **B)** Pam_2_CSK_4_. Median fluorescent intensity (MFI) ratio is determined by dividing the MFI of fluorescent ligand on CD4^+^ T cells by that of CD8^+^ T cells. Data represent mean ± s.d. (n = 3 independent samples).

**Table S1**: Quantitation of cell surface TLR2 ligands. Depoted Pam_2_CSK_4_ and Pam_3_CSK_4_ ligand concentrations calculated per T cell and per volume.

**Fig. S3**: B cells activate in presence of TLR2 ligand in a dose-dependent manner. **A)** Dose response of Pam_2_CSK_4_ and Pam_3_CSK_4_ in B-cell culture for 2 days as measured by fold MFI of B cells. Data represent mean ± s.d. (n = 3 independent samples). **B)** Dose response of Pam_2_CSK_4_ and Pam_3_CSK_4_ in B- and T-cell culture for 2 days as measured by fold MFI of B cells. Data represent mean ± s.d. (n = 2 independent samples).

**Fig. S4**: Viability of purified T cells after addition of IL-7. Cell viability of T cells that were **A)** in absence of (naïve T cells, left), or in presence of (primed T cells, right) 2 μg/ml of Concanavalin A and 10 ng/ml of IL-7 for 2 days and then resting in 10 ng/ml of IL-7 for up to 8 days. **B)** Cell viability of T cells that were without IL-7. Data depict m ± s.d (n = 1–2 independent samples).

**Fig. S5**: Depoted lipid-TLR2 ligand minimally enhanced proliferation indices of activated murine T cells. Purified polyclonal T cells were stained with 5 μM of carboxyfluorescein succinimidyl ester (CFSE). Different combinations of cell surface ligands (Pam_2_CSK_4_ and Pam_3_CSK_4_) were either directly added in solution (soluble) or depoted into polyclonal T cells for 1 h and cultured with αCD3/αCD28 beads for 3 days. Quantification of proliferation indices of **A)** CD4^+^ and **B)** CD8^+^ T cells in bulk polyclonal T cells as measured by CFSE dilution. Dotted lines represent respective averages (mean) of “No Ligand” conditions. p values between corresponding soluble vs depoted ligands as determined by one-tailed ratio paired t test; *p < 0.05; ns, not significant. Data depict m ± s.d. (n = 3 independent samples).

**Fig. S6**: Depoted lipid-TLR ligand T cells minimally enhanced proliferation of activated T cells. Purified polyclonal T cells were stained with 5 μM of carboxyfluorescein succinimidyl ester (CFSE). Different combinations of TLR2 ligands (Pam_2_CSK4 and Pam_3_CSK4) and TLR9 ligand (lipid-CpG) were either directly added in bulk solution (soluble) or depoted into polyclonal T cells for 1 h and cultured with αCD3/αCD28 beads for 3 days. **A)** Representative histograms of CD4^+^ T-cell proliferation from delivery of lipid-TLR ligand as measured by CFSE dilution. Quantification of division and proliferation indices of **B)** CD4^+^ and **C)** CD8^+^ T cells in bulk polyclonal T cells as measured by CFSE dilution. Dotted lines represent respective averages (mean) of “No Ligand” conditions. p values between corresponding soluble vs depoted ligands as determined by one-tailed ratio paired t test; *p < 0.05; ns, not significant. Data depict m ± s.d. (n = 3 independent samples).

**Fig. S7**: Lipid-TLR ligands promote promote Th1-based T-cell response. Different combinations of TLR2 ligands (Pam_2_CSK4 and Pam_3_CSK4) and TLR9 ligand (lipo-CpG) were either directly added in solution (soluble) or depoted into polyclonal T cells for 1 h and cultured with αCD3/αCD28 beads for 3 days. Quantification of **A)** IL-4 and **B)** IL-2 levels in T-cell supernatents as measured by ELISA. Dashed lines represent limit of detection for respective cytokine detection. (n = 2–3 independent samples).

**Fig. S8**: Lipid-TLR2 ligands depoted into murine T cells minimally enhance cell proliferation indices. Purified polyclonal T cells were stained with 5 μM of carboxyfluorescein succinimidyl ester (CFSE). Different combinations of TLR2 ligands (Pam_2_CSK_4_ and Pam_3_CSK_4_) were either directly added in bulk solution (soluble) or depoted into stained T cells for 1 h and cultured with non-depoted, stained T cells and αCD3/αCD28 beads for 3 days. Quantification of proliferation index of CD4^+^ and CD8^+^ T cells in bulk polyclonal T cells as measured by CFSE dilution. Data depict m ± s.d. (n = 5 independent samples).

**Fig. S9**: Depoted lipid-TLR ligand T cells increase CD25 expression on activated T cells. Purified polyclonal T cells were stained with 5 μM of carboxyfluorescein succinimidyl ester (CFSE). Different combinations of TLR2 ligands (Pam_2_CSK4 and Pam_3_CSK4) and TLR9 ligand (lipid-CpG) were either directly added in bulk solution (soluble) or depoted into polyclonal T cells for 1 h and cultured with αCD3/αCD28 beads for 3 days. CD25 expression as measured by MFI. p values between corresponding soluble vs depoted ligands as determined by two-way ANOVA with Sidak’s multiple comparisons test; *p < 0.05; ns, not significant. Data depict m ± s.d. (n = 3 independent samples).
